# Temporal matches and mismatches between monarch butterfly and milkweed population changes over the past 12,000 years

**DOI:** 10.1101/2022.02.25.481796

**Authors:** John H. Boyle, Susan Strickler, Alex Twyford, Angela Ricono, Adrian Powell, Jing Zhang, Hongxing Xu, Harmony J. Dalgleish, Georg Jander, Anurag A. Agrawal, Joshua R. Puzey

## Abstract

In intimate ecological interactions, the interdependency of species may result in correlated demographic histories. For species of conservation concern, understanding the long-term dynamics of such interactions may shed light on the drivers of population decline. Here we address the demographic history of the monarch butterfly, *Danaus plexippus*, and its dominant host plant, the common milkweed *Asclepias syriaca*, using broad-scale sampling and genomic inference. Because genetic resources for milkweed have lagged behind those for monarchs, we first release a chromosome-level genome assembly and annotation for common milkweed. Next, we show that despite its enormous geographic range across eastern North America, *A. syriaca* is best characterized as a single, roughly panmictic population. Using Approximate Bayesian Computation via Random Forests (ABC-RF), a machine learning method for reconstructing demographic histories, we show that both monarchs and milkweed experienced concurrent range expansion during the most recent recession of North American glaciers ∼12,000 years ago. Our data identify an expansion of milkweed during the large-scale clearing of eastern forests (∼200 years ago) but was inconclusive as to expansion or contraction of the monarch butterfly population during this time. Finally, our results indicate that neither species experienced a population contraction over the past 75 years. Thus, the well-documented decline of monarch abundance over the past 40 years is not visible in our genomic dataset, reflecting a possible mismatch of the overwintering census population to effective population size in this species.

## Introduction

Despite the critical importance of understanding past population dynamics, especially for species of conservation concern, inferring demographic histories can be extremely challenging. Novel genomic methodologies based on sampling extant individuals and interpretation of genomic patterns of diversity have recently provided insight into the demographic histories of species ranging from protists to humans (Schwabl et al. 2021; Lepers et al. 2021). Over the past 25 years, conservationists have become increasingly alarmed by the decline of the monarch butterfly’s overwintering population (Thogmartin et al. 2017; Pleasants et al. 2017; Lincoln P. Brower et al. 2012). Despite significant academic and public energy focused on understanding and reversing this, the exact cause of this decline is still a matter of debate. Multiple factors have been proposed to underlie the monarch’s decline, including a decrease in the abundance of the monarch’s food source (primarily a single species of milkweed – common milkweed), reduced abundance or quality of nectar plants, climate change, and destruction of their overwintering site (Haan and Landis 2019; Boyle, Dalgleish, and Puzey 2019; Inamine et al. 2016; Zylstra et al. 2021).

Here we address correlated demographic changes of monarchs and milkweeds over three hypothesized critical events during the Holocene. Placing this recent decline in a historical context will help us begin to address fundamental questions about the relationship between milkweed, monarchs, and humans. For instance, did colonizing Europeans inadvertently increase the size of the monarch population by massively expanding milkweed habitat through deforestation and ploughing of prairies? Does the recent decline of the overwintering census population follow from an artificial high? Or, does it represent a decline to levels lower than those seen before European colonization? And finally, are monarch and milkweed population demographics matched, perhaps indicating that milkweed is the limiting resource for monarch butterfly populations? Providing insight into these questions has remained intractable to date. However, recent advances in population genetic approaches and machine learning now allow us unprecedented ability to reconstruct demographic histories of populations. To reconstruct the demographic histories of monarchs and milkweed, here we use Approximate Bayesian Computation with Random Forests (ABC-RF) <u>(Pudlo et al. 2016)</u>. This approach has recently been employed by a number of population genetic studies on a diverse array of organisms, including humans <u>(Estoup et al. 2018)</u>, insects <u>(Lombaert et al., n.d.)</u>, plants <u>(Nevado et al. 2020)</u>, chordates <u>(Smith et al. 2018)</u>, and pathogens <u>(Schwabl et al. 2021)</u>, and it has been used to reconstruct biological invasions and other demographic events happening within the past few decades or centuries <u>(van Boheemen et al. 2017; Vallejo-Marín et</u> <u>al. 2021; Fraimout et al. 2017)</u>.

Accordingly, we use the ABC-RF approach to test how the last glacial retreat, the ploughing-up of the prairie and deforestation, and finally expansion of industrial agriculture impacted monarch and milkweed populations. Specifically, we addressed the following questions: (1) Have *A. syriaca* and *D. plexippus* populations expanded in prior millennia (12-5 kya), potentially due to the retreat of the glaciers after the last glacial maximum? (2) Have *A. syriaca* and *D. plexippus* populations expanded in the past centuries (1751-1899), potentially due to the conversion of native forests and prairies to agriculture land, as suggest by, e.g., Brower (1995) (L. P. Brower 1995)? (3) Have *A. syriaca* and *D. plexippus* populations experienced a bottleneck along with the industrialization of agriculture within past decades (1945-2015), potentially due to the increased use of herbicide in crop fields as described by, e.g., Pleasants (Pleasants and Oberhauser 2013, 2017)?

To facilitate answering these questions, we assembled a new genome for *A. syriaca.* Previously existing genomic resources are limited to low coverage assemblies and transcriptomes. Next, we sampled and conducted genomic analyses for 231 milkweed isolates from across the entire native range. Finally, using this data set, we test a series of explicit hypotheses using ABC-RF to ask how these climate and anthropogenic events have impacted population change of these iconic species. We conducted these analyses in parallel on milkweed and monarchs, using previously published whole-genome sequencing data from (Zhan et al. 2014) for the latter. As such, our analysis addresses whether the demographic histories of this intimate species interaction are matched or independent.

## Results

### Genome Assembly

Using PacBio and Hi-C libraries, we assembled a chromosome-level genome of 317 Mbp for *A. syriaca*, mostly assembled into 11 large chromosome-length molecules. A total of 42,111 genes were predicted, capturing 93% of the BUSCO set. Further details of this assembly are provided in the Supporting Information.

### SNP Calling

We gathered four different population genetic data sets for *D. plexippus* and *A. syriaca*:

For common milkweed:

(1) Broad Range WGR: We used a skimming Whole Genome Resequencing (WGR) approach at low coverage to identify approximately 900 SNPs from 48 plants collected from across the North American range of this species.
(2) Broad Range GBS: We used a Genotyping by Sequencing (GBS) approach to sequence and call approximately 900 SNPs from 96 plants collected from across the North American range of this species.
(3) Core Range GBS: We used a GBS approach to sequence and call approximately 900 SNPs from 87 plants, primarily collected in the eastern portion of this species’ range. This data set also includes a number of individuals collected from eastern Europe, where *A. syriaca* is an invasive species.

We analyzed the two different GBS datasets separately as they were produced in different labs and had different sequencing coverages.

For monarch butterflies, we used:

4) the whole genome sequences published by Zhan *et al*. (2014). From these we called approximately 11,700 Single Nucleotide Polymorphisms (SNPs) from 28 butterflies collected across the North American migratory range of this species.

We present the results from both species as parallel analyses. Sequencing localities for each milkweed data set are shown in Figure 1A, and more detailed results of the SNP calling process are provided in the Supporting Information.

**FIGURE 1:**
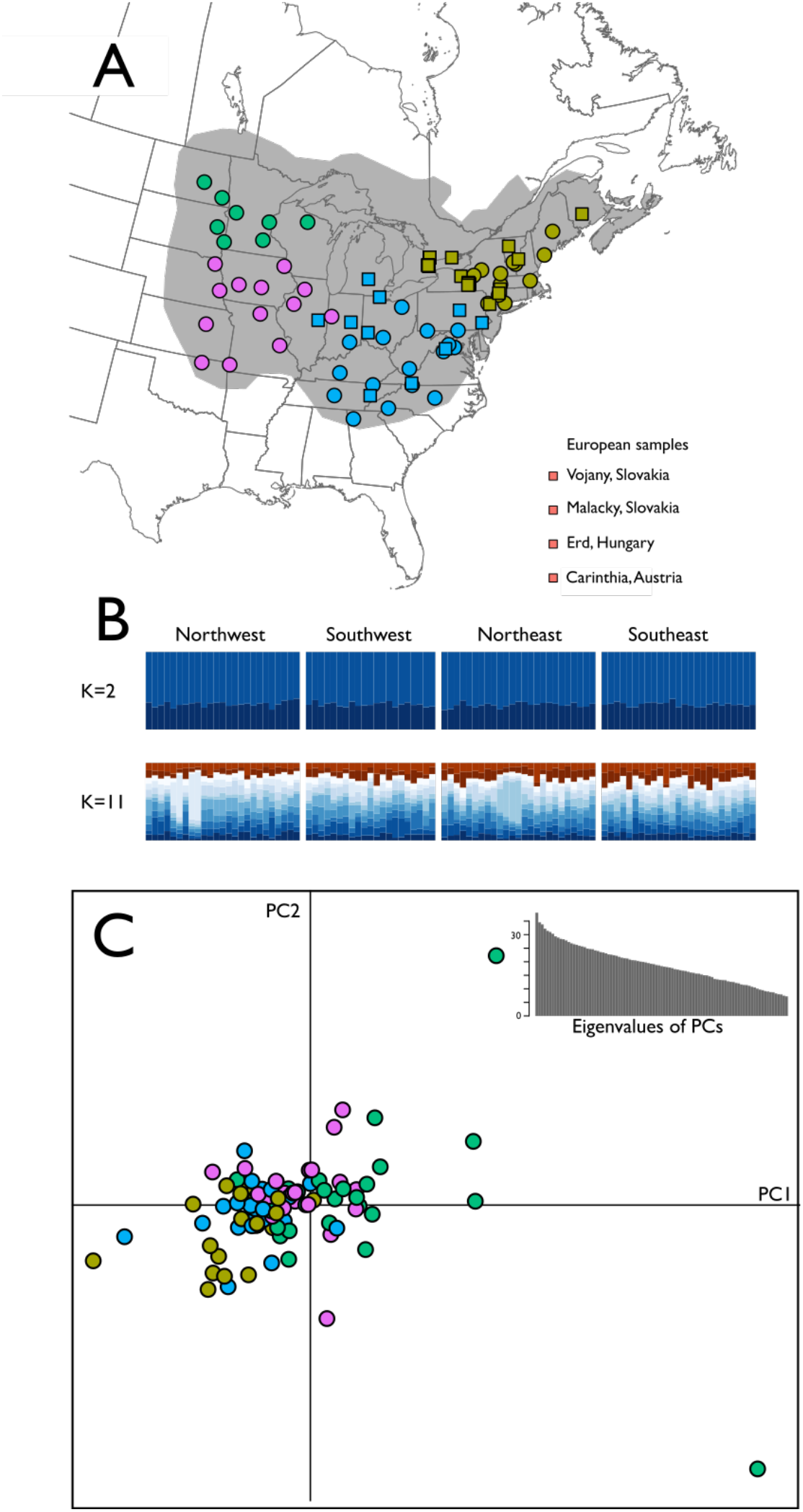
A: Our sampling scheme covers most of the North American range of *A. syriaca*. Circles represent sites sampled for the Broad Range data sets, while squares represent sites sampled for the Core Range data sets. Sites are colored according to the rough geographic zones to which we assigned them for the purposes of calculating Fst. We assigned the Core Range site in Illinois to the southeastern population instead of the southwestern population, since otherwise we would have only one locality representing a population in that data set. The gray region is an approximation of the range of *A. syriaca* based on specimen records in GBIF (“GBIF” 2021). B: STRUCTURE found no evidence of population structure among our milkweed specimens. The thin vertical bars represent individual milkweeds, and the four geographic zones are separated by thin white bars. Each bar is colored according to the cluster(s) to which it belongs. We present the results for the simplest analysis, in which STRUCTURE assumes K=2 clusters, and the analysis chosen by the Evanno method as optimal, K=11 (Evanno, Regnaut, and Goudet 2005). These results show strong genetic homogeneity across milkweed’s range. These data are from the Broad Range GBS data set; our other data sets produced similar results and are shown in the Supporting Information for all K-values from 2-20. C: PCA demonstrates weak geographic signal among some subsets of SNPs. Shown here are the first two principal components axes of allele frequencies, with each point representing an individual milkweed from the Broad Range GBS data set. Points are colored according to origin using the same color scheme as in Fig. 1A. The inset shows the eigenvalues for each principal component; these decline quite slowly, indicating that each individual PC axis explains relatively little of the variation in genotype. PC plots for additional axes, and for other data sets, show similarly weak levels of geographic signal, and are given in the Supporting Information.

### Population Genetic Analysis

All three of our milkweed data sets showed little genetic structure across their ranges. Global F_ST_ ranged from −0.002 (Broad Range WGR data set) to 0.039 (Core Range GBS data set), indicating a low amount of geographically sorted population structure. F_ST_ values between pairs of populations were similarly low, with the exception that the invasive European population was more distinct from the North American populations, with pairwise F_ST_ values around 0.08. We further interrogated this genetic structure using two approaches.

In the first approach, we used STRUCTURE to assign each individual ancestry to 2 or more subpopulations. It is important to note that STRUCTURE cannot be used to evaluate the fit of a single panmictic population as the optimal number of genetic clusters is determined based on the change in the log-likelihood between k-values [see (Janes et al. 2017)]. Regardless of the number of subpopulations chosen *a priori*, for every subpopulation, STRUCTURE assigned all individuals roughly the same degree of ancestry in that subpopulation, regardless of their geographic location (visualized in Figure 1B for the Broad Range GBS data set). This was true across all three data sets; the one major exception was that in the Core Range GBS data set, the invasive European population was quite distinct from the North American populations. STRUCTURE results for all three data sets are provided in the Supporting Information.

Secondly, to circumvent the inability of STRUCTURE to evaluate k=1, we took a less-parameterized approach by performing a Principal Components Analysis (PCA) on the allele frequencies of the SNPs in each data set. This approach identifies groups of covarying SNPs. We found a slight degree of geographic signal in several of the most important PC axes. For instance, in the Broad Range GBS data set (visualized in Figure 1C), PC1 largely separates several northwestern individuals from the remainder of the data set, possibly indicating introgression from *A. speciosa*, which is known to hybridize with *A. syriaca* in the northwestern part of the *A. syriaca* range. PC2 shows a slight amount of geographic signal, with western populations tending toward positive values and eastern populations tending toward negative values, but individuals from all four regions are well mixed in principal component space, indicating that this geographic signal is quite weak.

All three datasets support the conclusion that, in North America, *A. syriaca* is a single large metapopulation with little geographic structure. Our results parallel the findings of Zhan *et al*. (2014) that monarch butterflies, even between the eastern and western migratory populations, also lack geographic population genetic structure in North America. Additional information on each data set is given in the Supporting Information.

### Demographic modelling

We next used all four data sets (3 milkweed and 1 monarch) to estimate the recent demographic history of the two species. We used an Approximate Bayesian Computation (ABC) modelling approach, using a Random Forest (RF) algorithm for model selection.

Briefly, ABC modelling in population genetics uses simulated data sets to estimate posterior probabilities of past demographic events (Sisson, Fan, and Beaumont 2018), and the Random Forest approach described by Pudlo *et al*. (2016) (Raynal et al. 2019; Pudlo et al. 2016) which implements a machine learning algorithm to conduct the model selection. The ABC-RF approach allowed us to estimate posterior probabilities of each of the three events having occurred, separately for each data set. Additional details of these analyses may be found in the Supporting Information.

We found substantial consistency in the model predictions both across our three milkweed data sets and when comparing monarch and milkweed population histories. All three milkweed data sets show a post-glacial expansion with >0.9 posterior probability; indeed, this precisely matches the same strong prediction for monarchs (Figure 2).

**Figure 2:**
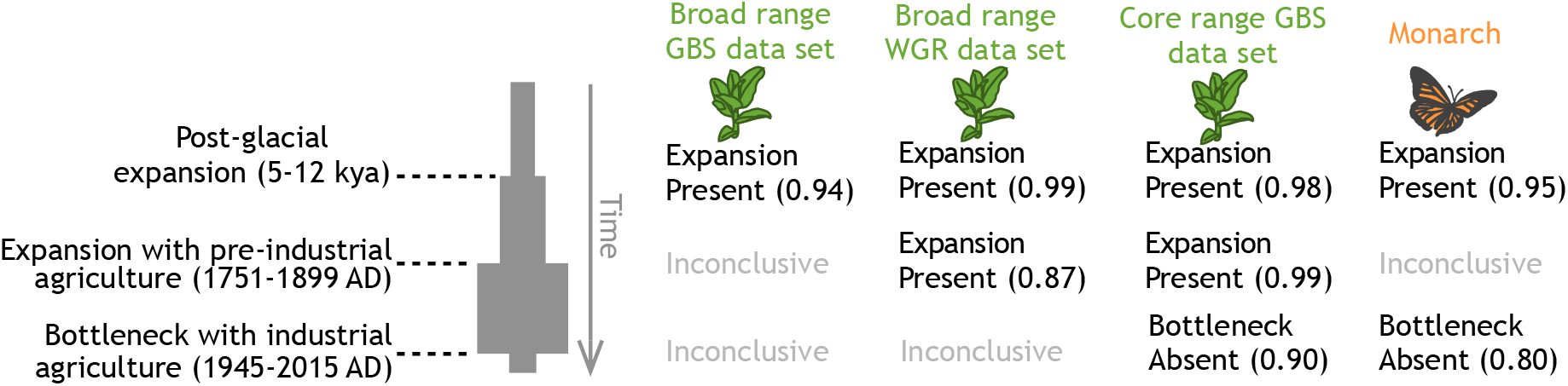
Support for each of our hypothesized demographic events in our three milkweed and one monarch data sets. The Random Forest consensus on whether each event is present in the population history of that species is given, along with the estimated posterior probability of each in parentheses. Posterior probabilities below 0.80 were considered “inconclusive”; posterior probabilities for all demographic events are given in Table S2.

In milkweed, we found evidence for a centuries-scale expansion alongside agriculture in the 18^th^ and 19^th^ centuries. The Core Range GBS and Broad Range WGR data sets showed strong support for the presence of such a population expansion (0.99 and 0.87 posterior probabilities, respectively). The Broad Range GBS data set was inconclusive (posterior probability of 0.61 that such an expansion did not exist). We consider the Broad Range GBS dataset as inconclusive as it did not have enough evidence to shift the posterior probability far from the prior probability of 0.50. The monarch data were also inconclusive on the presence of an expansion during this period (0.68 posterior probability in favor of an expansion).

The absence of a recent bottleneck alongside the industrialization of agriculture was weakly supported in the monarch data set (0.80) and in the Core Range GBS milkweed data set (0.90), but the other two milkweed data sets were inconclusive, with no support for either the presence or the absence of a recent bottleneck (Figure 2).

## Discussion

Understanding the impact of the Anthropocene on the natural world is of fundamental importance for conservation efforts. Until recently, elucidating patterns of population change in the recent past has been very difficult. In this study we employ an ABC-RF approach to study the near-term demographic history of monarchs and milkweeds. This approach was chosen in part because it has proven useful in other systems in elucidating very recent demographic events, within decades or centuries (Vallejo-Marín et al. 2021; Fraimout et al. 2017). In addition, this approach requires fewer simulated datasets to train the classifier than are necessary for traditional ABC, and it is much more robust to choices of summary statistics (Pudlo et al. 2016; Csilléry, François, and Blum 2012).

We tested for changes in effective population size of the monarch butterfly and its primary food source, common milkweed, during three events: the most recent retreat of the glaciers, European settlement, and industrial agriculture. Previously, using PSMC (Pairwise Sequentially Markovian Coalescent) model, a method capable of testing for ancient events but less fit for resolving recent events, researchers demonstrated a population expansion of monarch butterflies after the last glaciation (Zhan et al. 2014). Using ABC-RF, we likewise detect this monarch expansion and also observe an expansion of common milkweed post-glaciation. The low levels of population structure in common milkweed likely occur because the modern range of *A. syriaca* is a result of rapid (i.e., in the last 5-12 kya) invasion of central and eastern North America after the retreat of the glaciers. In this scenario, the rapid expansion combined with *A. syriaca* being an obligate outcrosser with long-distance dispersal ability, has prevented the formation of extensive population structure.

We provide population genetic evidence that common milkweed increased in prevalence during the 18th and 19th centuries. The most obvious cause for this is the clearing of forests and prairies to make way for agricultural land, a disturbance-rich environment in which *A. syriaca* thrives (at least, until the advent of herbicides). The increase observed in our data has previously been suspected, and there are two major hypotheses for how this increase affected monarch butterflies. The first hypothesis posits that *A. syriaca* has always been the most important host plant for monarchs, even before *A. syriaca*’s population boom. As *A. syriaca* increased in abundance in a newly-disturbed landscape, monarchs increased in abundance alongside them. Thus, according to this hypothesis, the current size (and possible geographic extent) of the monarch migration was greater in the 18th-20th centuries than in the 17th century and prior (Brower 1995; a more radical form of this hypothesis suggests that the migratory behavior itself was absent before the 18th century, Vane-Wright 1993) (L. P. Brower 1995; Vane-Wright 1993). However, although *A. syriaca* has increased in abundance due to disturbance, it is likely that other species of milkweeds, less tolerant of anthropogenic changes, have declined in abundance over the same period. The second hypothesis suggests that monarch transitioned from a wider array of host plant species to become more reliant on common milkweed over this period of increase in common milkweed populations. If this occurred, then the newly-increased population sizes of *A. syriaca* did not represent a net increase in food resources for monarchs, and so we would not expect the monarch abundances in the 18th-20th centuries to be higher (or lower) than previously (Brower 1995).

How should biologists and conservationists react to this new data that shows common milkweed increased with agriculture? This depends largely on which hypothesis about the monarch response to this increase is correct. If the 20th century population size of the monarch was anthropogenically inflated due to elevated common milkweed abundance, this puts contemporary declines in a less worrisome light, as they may simply represent returns to pre-modern population sizes. Monarch population sizes and migratory behavior have presumably been sustainable for centuries before the clearing of the forests and prairies of Eastern North America. However, if monarchs responded to increased common milkweed abundance by shifting their diets without increasing the total population, then contemporary declines may well have put the monarchs at their lowest population size since the retreat of the glaciers.

Unfortunately, the whole-genome-sequencing data set used here was not sensitive enough to detect whether or not there had been an increase in monarch populations alongside those of milkweed in the 18th and 19th centuries. Answering this question using population genetics will probably require improvements in our current techniques for demographic modelling and/or denser sequencing of *D. plexippus* individuals than is currently available. However, there are other potential data sets that could shine light on this question. As a start, population genomic analyses for other important milkweed species could reveal whether or not they declined during the period of common milkweed’s increase: lack of such declines would suggest that the expansion of *A. syriaca* in particular could only have increased the monarch population. Brower (1995) suggests sampling cardenolide profiles from museum specimens of monarchs captured in the 19th and 20th centuries. These profiles can indicate the host plants those individuals used as larvae, and thus show whether or not monarchs experienced a shift in their host species as humans cleared forests and prairies. Shifts to more diversity in milkweed hosts might also be detectable in more recent specimens collected on the East Coast of North America, as farming has become less prevalent in this region over past decades: the presence or absence of such shifts would be evidence that the opposite had happened when this same region of the country was being deforested in the 18th and 19th centuries.

Finally, even if such tests demonstrate that the current declines in monarch butterfly populations are simply a return to historically-sustainable population sizes, we emphasize that this is not necessarily an argument against current efforts to support monarch butterfly populations. Regardless of how many monarchs were in North America in 1600, the current monarch population brings delight to people across North America and serves as a key conservation species which serves as an introduction to many non-scientists to the importance of invertebrate conservation, pollination biology, migratory behavior, and more. Having fewer of these charismatic insects present would be a loss to humankind regardless of how many of them were present a few centuries ago.

### Abbreviated Methods

Additional details of all the methods described below are given in the Supporting Information.

### Genome assembly

Genomic DNA was prepared from one individual of *Asclepias syriaca* collected from Austria (46.66,14.47) and sequenced using PacBio CLR technology on six SMRT cells. Illumina sequence was generated from genomic DNA on one lane of Hi-Seq 2 x 150 bp. Hi-C libraries were prepared using the Proximo Hi-C kit for plants (Phase Genomics) and sequenced on one lane of Illumina NexSeq500 2 x 150 bp. The genome was assembled from these data, haplotigs purged, and scaffolding performed as described in the Supporting Information.

### SNP calling

Two sets of GBS libraries and one set of WGR libraries were prepared as described in the Supporting Information. SNPs from the GBS dataset were identified using Stacks 2.2 (Catchen et al. 2013; Rochette, Rivera-Colón, and Catchen 2019). SNPs from the WGR data set were called using the Genome Analysis Toolkit (GATK) pipeline (McKenna et al. 2010; DePristo et al. 2011). We also used the GATK pipeline to call SNPs from the sequence data provided in Zhan *et al*. (2014) for a number of North American monarchs (Zhan et al. 2014).

### Population genetic analysis

We assigned each individual milkweed to one of 5 broad geographic populations based on its location, as shown in Figure 1A. We used the hierfstat package (Goudet 2005) in R to calculate F_ST_ for each data set. To examine clustering and admixture within the *A. syriaca* populations, we used STRUCTURE 2.3.4 (Pritchard, Stephens, and Donnelly 2000), using and an admixture model. Finally, we used the ade4 package (Daniel Chessel Anne B. Dufour Jean Thioulouse 2004) in R to perform a PCA on the allele frequencies for each of the milkweed data sets.

### Demographic modelling

To investigate the recent demographic history of *A. syriaca*, we used an Approximate Bayesian Computation (ABC) modelling approach for model selection. We used the Random Forest approach described by Pudlo *et al*. (Pudlo et al. 2016), which implements a machine learning algorithm to do model selection. As our observed data, we used separately each of the four monarch or milkweed data sets produced above. Guided by the results of our STRUCTURE analysis, we treated *A. syriaca* as a single population. We simulated data sets using DIYABC 2.1.0 (Cornuet et al. 2014) to test the three hypotheses visualized in Figure 2.

We then constructed a random forest of 1000 decision trees, each of which provided a prediction of which demographic model produced a given set of summary statistics. We then fed our observed data set into this random forest in order to estimate the best model and approximate its posterior probability.

### Data Availability

Data and scripts will be made publicly available on Dryad and Genbank upon acceptance (Genbank Project ID: PRJNA787127).

## Contributions

**T**his project was conceived of by JB, GJ, AA, and JP. The assembly and annotation of the milkweed genome was performed by SS, AP, JZ, GJ, and HX. Collection of milkweed samples, DNA extractions, and DNA library preparation was done by AR, HD, HX, and AT. Design of ABC portion of the project was overseen by JB and JP. JB conducted the population genetic analyses. The paper was primarily written by JB and JP with significant editing input from AA and AT. All authors approved the final text.

## Acknowledgments

This work was supported by NSF 1645256 (Jander and Agrawal), USDA 2020-67013-30896 (Jander), Triad Foundation (Jander), Jeffress Trust Awards Program in Interdisciplinary Research (Puzey), Dominion Education Partnership (Puzey and Dalgleish), and National Geographic GR-000000959 (Puzey and Dalgleish).

## Supporting Information

### MATERIALS AND METHODS

#### Milkweed Husbandry

Common milkweed (*Asclepias syriaca*) seeds were sterilized in 3% sodium hypochlorite containing 0.05% Tween 20 for 10 min and rinsed with sterile water for 5 times. After being scarified and cold-stratified at 4 °C on moist filter paper for 2 weeks, the seeds were germinated in a dark warm chamber at 28 °C for 4-5 days. The seedlings were planted into potting soil (60% lamberts, 20% perlite and 20% turface) (9-cm square pots) and grown completely randomized in a growth chamber under a 16:8 h day: night cycle at 23 °C with a 60% relative humidity. Older plants were moved to a greenhouse with natural sunlight.

#### Genome assembly

##### Genome sequencing and assembly of A. syriaca

Genomic DNA was prepared from one individual of *Asclepias syriaca* and sequenced using PacBio CLR technology on six SMRT cells. Illumina sequence was generated from genomic DNA on one lane of Hi-Seq 2 x 150 bp. Kmer analysis was performed using this Illumina sequence, Jellyfish(Marçais and Kingsford 2011), and Genomescope(Vurture et al. 2017). Hi-C libraries were prepared using the Proximo Hi-C kit for plants (Phase Genomics) and sequenced on one lane of Illumina 2 x 150 bp. *A. syriaca* PacBio sequence was assembled using Falcon v 2017.11.02-16.04 and falcon-kit 1.3.0(Chin et al. 2016) and the configuration file (<u>fc_run.cfg</u>) (Chin et al. 2016). The assembly was corrected using the Illumina sequence and Pilon v1.23. Redundancy was removed using Purge Hapolotigs (Roach, Schmidt, and Borneman 2018). Hi-C was used to scaffold the contigs using 3D-DNA v 180419 (Dudchenko et al. 2017) and gaps were filled with LR_gapcloser (G.-C. Xu et al. 2019) and corrected PacBio reads.

##### Genome annotation of A. syriaca

For repeat identification and masking, LTR_retriever (Ou and Jiang 2018) was used with outputs from LTRharvest (Ellinghaus, Kurtz, and Willhoeft 2008) and LTR_FINDER (Z. Xu and Wang 2007) to identify long terminal repeat retrotransposons (LTRs). The LTR library was then used to hard mask the genome, and RepeatModeler version: open-1.0.11 (Smit, AFA, Hubley, R & Green, P., n.d.) was used to identify additional repetitive elements in the remaining unmasked segments of the genome. Protein-coding sequences were excluded using blastx v2.7.1+ (Ellinghaus, Kurtz, and Willhoeft 2008; Altschul et al. 1990) results in conjunction with the ProtExcluder.pl script from the ProtExcluder v1.2 package (Campbell et al. 2014). The libraries from RepeatModeler and LTR_retriever were then combined and used with RepeatMasker version: open-4.0.7 (Smit, AFA, Hubley, R & Green, P., n.d.) to produce the final masked version of the genome.

Libraries with an insert size of 350 bp were prepared from leaf RNA and sequenced on one lane of 2 x 100 bp Illumina Hi-Seq. RNA-seq reads were mapped to the genome with HISAT2 v2.2.0 (Kim, Langmead, and Salzberg 2015). Portcullis v 1.1.2 (Mapleson et al. 2018) and Mikado v 1.2.2 (Venturini et al. 2018) were used to process and filter the resulting bam files. Augustus v 3.2.0 (Stanke et al. 2008) and Snap v 2006-07-28 (Korf 2004) were trained and implemented through the Maker v 2.31.10 pipeline (Cantarel et al. 2008), with proteins from Swiss-Prot (Boutet et al. 2007) and processed RNA-seq added as evidence. Gene models were filtered with the following criteria: 1) at least one match found in the Trembl database (4-17-19) (Boutet et al. 2007) with an E-value less than 1e-20, 2) InterProScan matches to repeats were removed, 3) genes with an AED score of 1 and no InterPro domain were removed, and 4) single-exon genes with no InterPro domain were removed. Functional annotation and classification were performed using BLASTx v2.7.1+ (Altschul et al. 1990) and InterProScan v5.36-75.0 (Jones et al. 2014). Both genome and annotation completeness were assessed by BUSCO v3.1.0 (Waterhouse et al. 2017) using the embryophyta lineage.

#### SNP Calling

##### Genotyping by sequencing (GBS) of the A. syriaca Core Range data set

A total of 283 common milkweed plants collected from different places around US and Europe were germinated and cultivated in our greenhouse. Fresh collected tissue was flash frozen in liquid nitrogen. The DNA was extracted from the leaf of individuals using a CTAB (cetyltrimethyl ammonium bromide)-based extraction protocol (adapted from <u>(Fulton,</u> <u>Chunwongse, and Tanksley 1995)</u>. The DNA was quantified using a CFX384 C1000 Real-Time thermal cycler (BioRad, Hercules, CA) and normalized to 30–100 ng/ul using a GBFit Arise Pipetting System (Pacgen Inc., Irvine, CA). Quality checks were performed by agarose gel observation of 300 ng of undigested and *Hind*III digested DNA per sample. Genotyping was performed following the GBS protocol <u>(Elshire et al. 2011)</u>, using *Ape*KI as the restriction enzyme. The libraries were sequenced on a HiSeq 2500 system (Illumina Inc., USA) with the single-end mode and read length of 101 bp.

##### Genotyping by sequencing (GBS) of the A. syriaca Broad Range data set

DNA was extracted from flash-frozen leaf samples using the Qiagen DNeasy Plant extraction kit. 100ng of sample DNA was used for GBS library preparation using the ApeKI restriction enzyme, as above. 95 samples and a water control (blank) were pooled per multiplex and sequenced using 100bp single-end mode on the HiSeq 2500 at the University of Rochester Medical Center.

##### Whole Genome Resequencing (WGR) of the A. syriaca Broad Range data set

DNA was extracted from *A. syriaca* using Qiagen DNeasy kit and sequenced using Illumina HiSeq 2×150.

##### SNP calling of the A. syriaca Core and Broad Range GBS data sets

Genotyping By Sequencing reads were demultiplexed using Stacks 2.2 (Rochette, Rivera-Colón, and Catchen 2019; Catchen et al. 2013). Reads from each individual where then mapped against the *A. syriaca* genome using Bowtie2 2.3.2 (Langmead and Salzberg 2012), using end-to-end alignment and the “--very-sensitive” alignment settings. Reads with a mapping quality lower than 5 were discarded using samtools 1.5 (Li et al. 2009). We then used Stacks in combination with custom scripts to call SNPs and to filter low-quality individuals and loci from our data set. The scripts will be deposited upon acceptance to Dryad. Briefly, several individuals in our data set had been identified as possible *A. speciosa* or *A. syriaca x A. speciosa* hybrids. Since *A. syriaca* and *A. speciosa* can be difficult to distinguish when they are not in flower, we did an initial clustering of our data using the find.clusters function implemented in adegenet 2.1.1 (Jombart 2008; Jombart and Ahmed 2011) in R 3.5.2 (R Core Team 2018). This identified several more putative *A. speciosa* individuals, which were removed.

Since *A. syriaca* can reproduce asexually, we also screened our data set for clones; i.e., different ramets of the same genet. To do so, we considered all pairs of individuals, calculating what percentage of their homozygous loci had identical SNP calls. Across all pairs of individuals, this distribution was bimodal. The vast majority of pairs were normally distributed around a sequence identity of 0.936, with a small number of comparisons clearly outside of this distribution, clustered around 1.00. Accordingly, we considered all pairs of individuals with a sequence identity greater than 0.993 to be clones. Where clones were found at the same site, we randomly selected a single exemplar, discarding all its clones from the data set. A few pairs of clones were found in different sites; in this case we discarded both members of the pair.

Combining the Broad Range and Core Range GBS Data Sets in subsequent analyses produced strong batch effects between the two data sets (see below), likely because they were sequenced on different machines, at different times, to different read depths. We therefore performed the following analyses separately for the two data sets.

After discarding *A. speciosa*, clones, and individuals for which relatively few loci (i.e., less than 80% of the total number of loci) had been sequenced, we then randomly downsampled the Core Range data set to include a maximum of 5 individuals per site, to homogenize sampling effort across the sites. Finally, we used Stacks to filter SNPs across these individuals, including SNPs with observed heterozygosity less than or equal to 0.6 and present in at least 80% of individuals. Where multiple SNPs were found at the same GBS locus, we randomly excluded all but one. To reduce linkage disequilibrium, we filtered SNPs so that each was at least 50 kb from its nearest neighbor.

We also used this data set, after excluding invasive individuals collected from Europe using vcftools 0.1.15 (Danecek et al. 2011), for demographic modelling. This data set was converted to DIYABC format using vcf2diyabc.py (“DIYABC” 2015).

##### SNP Calling of the A. syriaca Broad Range WGR data set

We called SNPs using the Genome Analysis Toolkit (GATK) pipeline (McKenna et al. 2010; DePristo et al. 2011; Van der Auwera et al. 2013). Reads from each individual were mapped against the *A. syriaca* genome using Bowtie2 2.3.2, with an expected range of inter-mate-pair distances of 100-2000 and the “--very-sensitive-local” alignment settings. Indicies of the genome were first built using both bowtie2 and samtools, and a sequence dictionary created using Picard 2.18.15 from the Genome Analysis Toolkit (Van der Auwera et al. 2013; McKenna et al. 2010; DePristo et al. 2011).

We further used Picard to fix mate pair information, mark and remove duplicate reads, and replace read group names; we then used samtools to index the alignments for each resequenced individual. We then called polymorphisms for each individual with the HaplotypeCaller tool, then combined the outputs from each scaffold using GenomicsDBImport. We then used GenotypeGVCFs to do joint genotyping on all individuals simultaneously. Indels were removed with the SelectVariants tool, and the remaining SNPs were filtered using the VariantFiltration tool, discarding SNPs for which any of the following were true: quality by depth (QD) less than 2; phred-scaled p-value of Fisher’s Exact Test for strand bias (FS) greater than 60; root mean square of the mapping quality (MQ) less than 35; mapping quality rank sum test (MQRankSum) less than -12.5; read position rank sum test (ReadPosRankSum) less than -8. We also filtered out loci with greater than 5% missing data or a minimum read depth of less than 5, as well as removing individual genotypes with a minimum quality 5 or less. Finally, SNPs were thinned to be 50 kb apart or more, so as to match the amount of thinning done for the GBS data set.

##### SNP Calling of the D. plexippus WGR data set

We used the whole genome sequencing data of Zhan *et al*. (2014) to gather genomic data from 29 monarch butterflies collected in North America (which individual specimens we used are given in Supplementary File SA; we chose migratory individuals from the continental United States and Mexico, excluding non-migratory individuals from South Florida) (Zhan et al. 2014). We called SNPs using the pipeline described above, aligning reads from each individual to the *D. plexippus* genome of Zhan *et al*. (2011), GenBank accession GCA_000235995.2 (Zhan et al. 2011). SNPs were filtered using the same criteria as for the *A. syriaca* WGR data, except that SNPs were thinned to be 10 Mb apart or more in order to produce a similar number of SNPs to those found in the *A. syriaca* data sets.

#### Population Genetic Analysis

##### F_ST_ analysis and basic population genetic statistics

Using all three *A. syriaca* data sets, and the *D. plexippus* data set, we estimated several population genetic statistics in R, using the adegenet and hierfstat packages (Goudet 2005; Paradis et al. 2017). We assigned each individual to one of five broad geographic populations based on its location. Population assignments are shown in Figure 1A. We tested whether this arrangement captured significant genetic structuring using an AMOVA test, using the pegas method (Paradis 2010) as implemented in poppr 2.8.2 (Kamvar, Tabima, and Grünwald 2014) with 10,000 permutations.

##### STRUCTURE analysis

To examine clustering and admixture within the *A. syriaca* populations, we used STRUCTURE 2.3.4 (Pritchard, Stephens, and Donnelly 2000). We analyzed all three data sets using an admixture model within STRUCTURE and all possible values for the number of clusters (*k*) between 1 and 20; running 10 replicates for *k* value. We chose the best number of clusters using the Evanno method (Evanno, Regnaut, and Goudet 2005) as implemented in Structure Harvester 0.6.94 (Earl and vonHoldt 2012). We also used Structure Harvester to convert STRUCTURE output files for use with CLUMPP 1.1.2 (Jakobsson and Rosenberg 2007). We used CLUMPP to assign consistent cluster identities across our multiple replicates for each *k* value above 1, using the LargeKGreedy algorithm with 1000 random input orders and the G’ matrix similarity statistic.

##### PCA analysis

To complement our STRUCTURE analysis, we also performed a PCA analysis to examine geographic distribution of genetic structure in a less parameterized way using the ade4 (Daniel Chessel Anne B. Dufour Jean Thioulouse 2004; Dray and Dufour 2007) and adegenet (Jombart 2008; Jombart and Ahmed 2011) packages in R. We first scaled each genotype using the scaleGen() function, replacing missing data with the mean allele frequency for that SNP, and then performed a Principle Components Analysis on these scaled allele frequencies.

#### Demographic modelling

To investigate the recent demographic history of monarchs and common milkweed, we used an Approximate Bayesian Computation (ABC) modelling approach, using a Random Forest (RF) algorithm for model selection and parameter estimation. Briefly, ABC modelling uses simulated data sets to estimate posterior probabilities when the likelihoods of observed data given specific models are difficult to calculate (Sisson, Fan, and Beaumont 2018; Beaumont, Zhang, and Balding 2002). Genetic data sets are simulated under a number of different demographic models, and the simulated data sets closest to the observed data are used to estimate the posterior probabilities of individual models and distributions of parameters of interest. We used the Random Forest approach described by Pudlo *et al*. and Raynal *et al*., which implements a machine learning algorithm to do model selection and parameter estimation (Raynal et al. 2019; Pudlo et al. 2016). The RF approach improves upon traditional ABC modelling in that ABC-RF is insensitive to the choice of summary statistics, and less computationally expensive as well.

As our observed data, we used the four monarch and milkweed data sets described above. Guided by the results of our STRUCTURE analysis, we treated *A. syriaca* as a single population. We simulated data sets using DIYABC 2.1.0 (Cornuet et al. 2014) to test the following hypotheses:

1. Have *A. syriaca* populations experienced a bottleneck within past decades, potentially due to the increased use of herbicide in crop fields as described by, e.g., Pleasants (2017)?
2. Have *A. syriaca* populations expanded in the past centuries, potentially due to the conversion of native forests and prairies to agriculture land, as suggest by, e.g., Brower (1995)?
3. Have *A. syriaca* populations expanded in prior millenia, potentially due to the retreat of the glaciers after the last glacial maximum (CITE)?

Considering every possible combination of the three hypotheses produced 8 demographic scenarios. We used DIYABC to simulate 80,000 data sets across all 8 demographic scenarios. For each scenario, population sizes were selected from uninformative prior distributions (see Table Da for details), while event times were chosen from uniform distributions. We chose event times to correspond to 1945-2015 for the recent bottleneck, 1751-1899 for the recent expansion, and 5-12 thousand years ago for the ancient expansion. *A. syriaca* plants flower in their second growing season (Bhowmik and Bandeen 1976), so we assumed a 2 year generation time for this species. *D. plexippus* has 4-5 generations per year, so we assumed a 0.2-0.25 year generation time for that species, which produces the values shown in Table Db. We outputted all 4 summary statistics calculated by DIYABC, which would be used for ABC-RF model selection, alongside the linear discriminant axes that were the combinations of those summary statistics that best distinguished the demographic models.

Following Pudlo *et al*. (2016), and using the abcrf package in R, we performed a number of validations of our ABC-RF approach: We first tested the compatibility of our models with our observed data by projecting our observed data, along with the simulations, along the linear discriminant (LD) axes that best distinguished the 8 models given the set of 4 summary statistics (Pudlo et al. 2016; Raynal et al. 2019). We then constructed a random forest of 1000 decision trees, each of which provided a prediction of which demographic model produced a given set of summary statistics. To test whether we had produced a sufficient number of simulations, we compared the prior error rate of this random forest to that of a second random forest constructed using only 80% of the 80,000 simulations. Finally, to test whether 1000 decision trees was a sufficient number, we calculated the prior error rate using forests of different size, from 40-1000.

We then fed our observed data set into this random forest in order to estimate the best model and approximate its posterior probability. Because the posterior probability of any single model was low, we produced separate random forests to approximate posterior probabilities for each of the three hypotheses listed above, i.e., by grouping together all models that had a recent bottleneck vs all models that did not, etc.

We then used the approach of Raynal *et al*. (2019), employing the ABC-RF approach to estimate parameter values (Raynal et al. 2019). We first used DIYABC to simulate 10,000 data sets for the single best demographic scenario. We then used this simulation set to estimate posterior medians and quantiles of a number of demographic parameters using ABC-RF.

### RESULTS

#### Genome assembly

##### Genome sequencing and assembly of A. syriaca

PacBio sequencing resulted in over 300X coverage of the expected genome size of 420 Mb[1]. The sequence was assembled into 748 contigs with a total length of 362 Mbp and an N50 of 1.9 Mbp. Kmer analysis supports this genome size. After haplotig removal, approximately 91% of the sequence was scaffolded into 11 sequences representing pseudomolecules. The final assembly has a length of 317 Mbp and captures 96.8% of the BUSCO set.

##### Genome annotation of A. syriaca

Approximately 57% of the genome consists of repetitive sequences. A total of 42,111 genes were predicted with an average length of 2,578 bp. Approximately 93% of the BUSCO protein set was identified in the annotation. Putative functions were assigned to 99% of the gene set.

#### SNP Calling

Collection sites and sample sizes for each data set are shown in Figure 2. The number of individuals and loci, and the amount of missing data for each SNP data set is shown in Table 1.

**Figure SI1:**
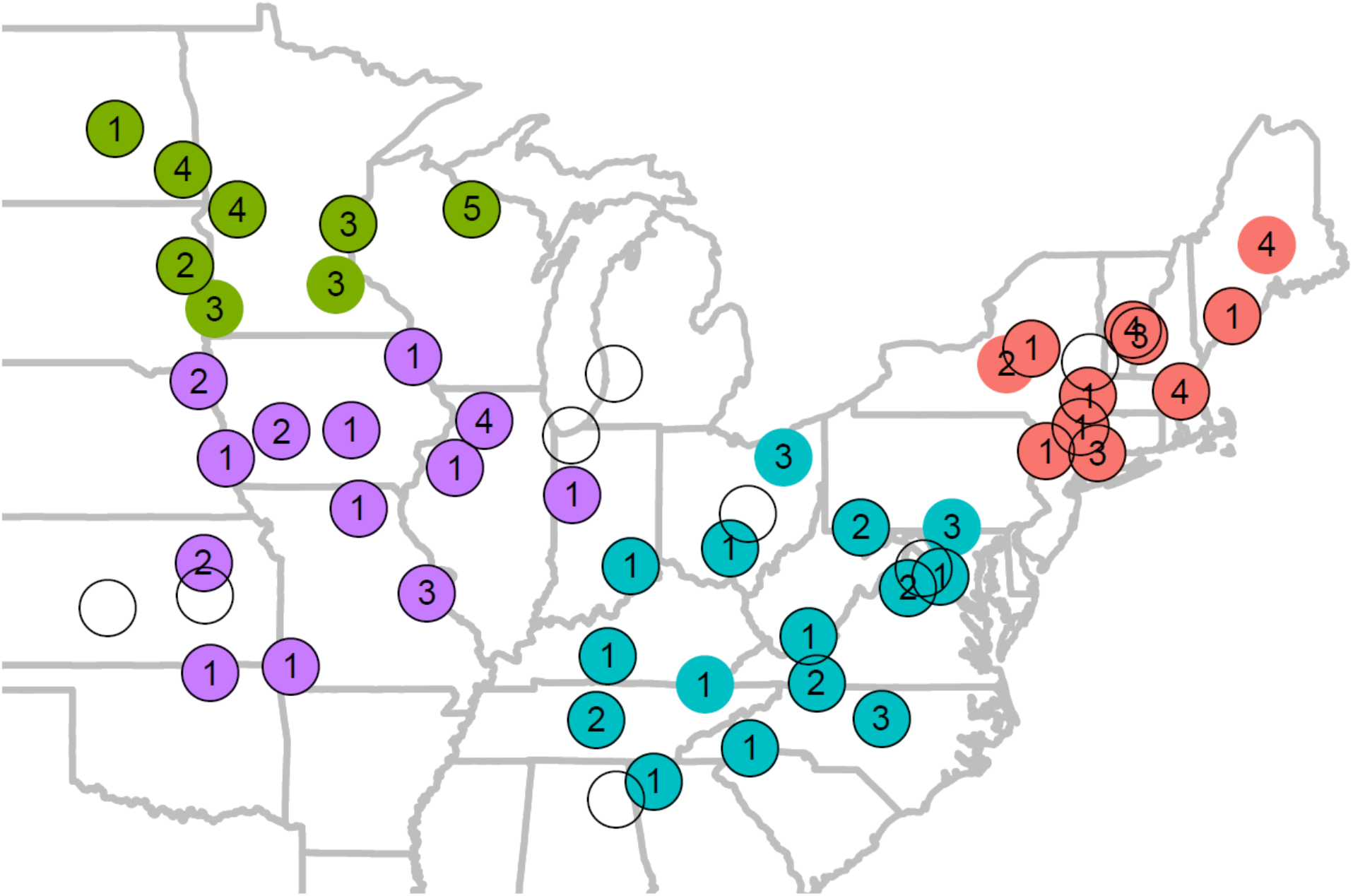
Sample scheme for the Broad Range milkweed data sets. Filled circles show collection localities for the Broad Range GBS data set; the number inside the circle shows the number of samples from each site included in this data set. Sites with a black border around the circle indicate that one individual from that location was used in the Broad Range WGR data set. Hollow circles indicate that one individual from that locality was used for the Broad Range WGR data set only. Sites are colored according to the population to which they were assigned: Green = Northwest, Purple = Southwest, Teal = Southeast, Red = Northeast.

**Figure SI2:**
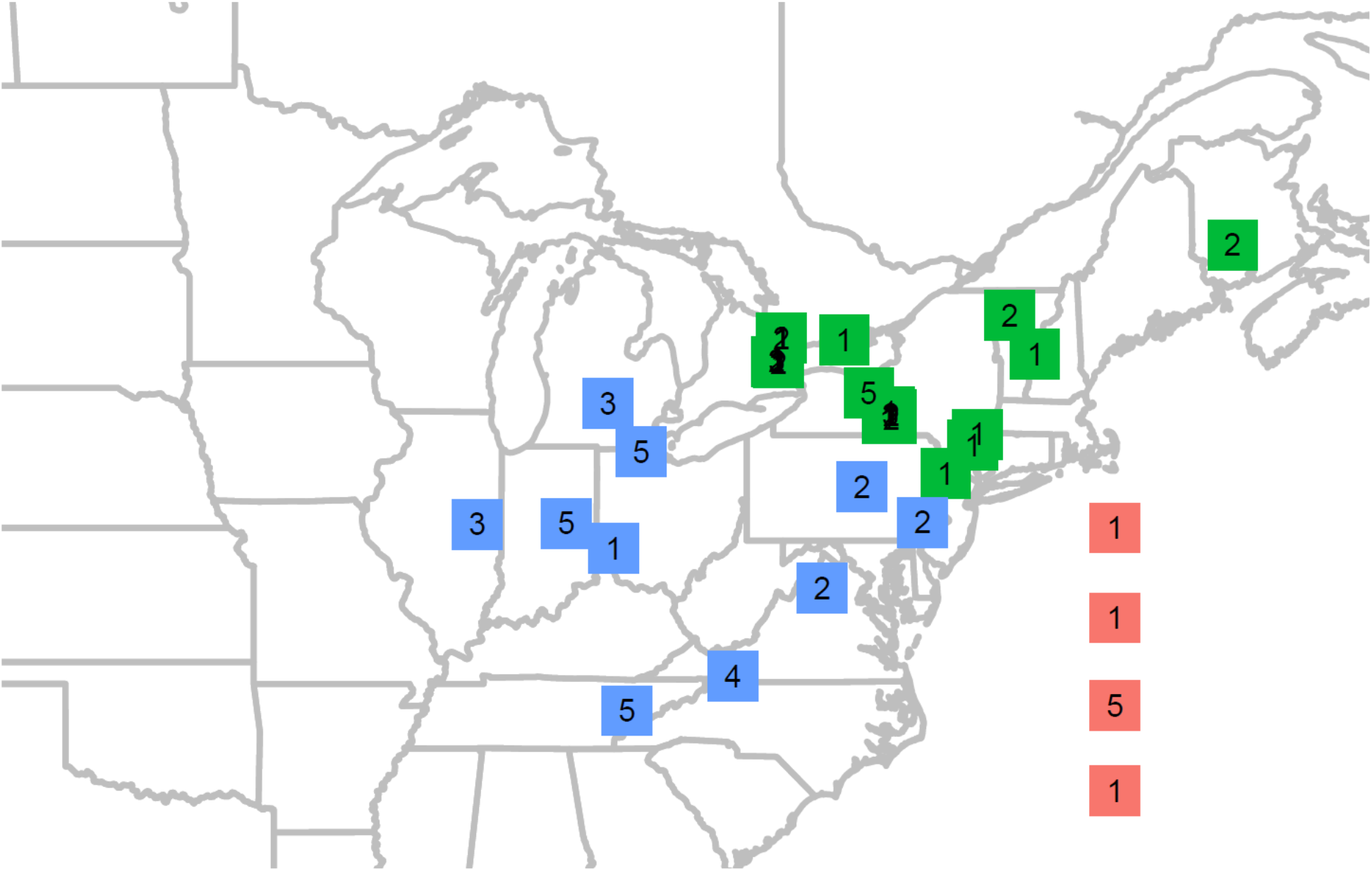
Sampling scheme for the Core Range milkweed data set. Squares show collection localities; the number inside the square shows the number of samples from each site included in the population genetic data set. Squares in the Atlantic ocean represent European collection sites. Sites are colored according to the population to which they were assigned: Green = Northeast, Blue = Southeast, Red = Europe.

**Figure SI3:**
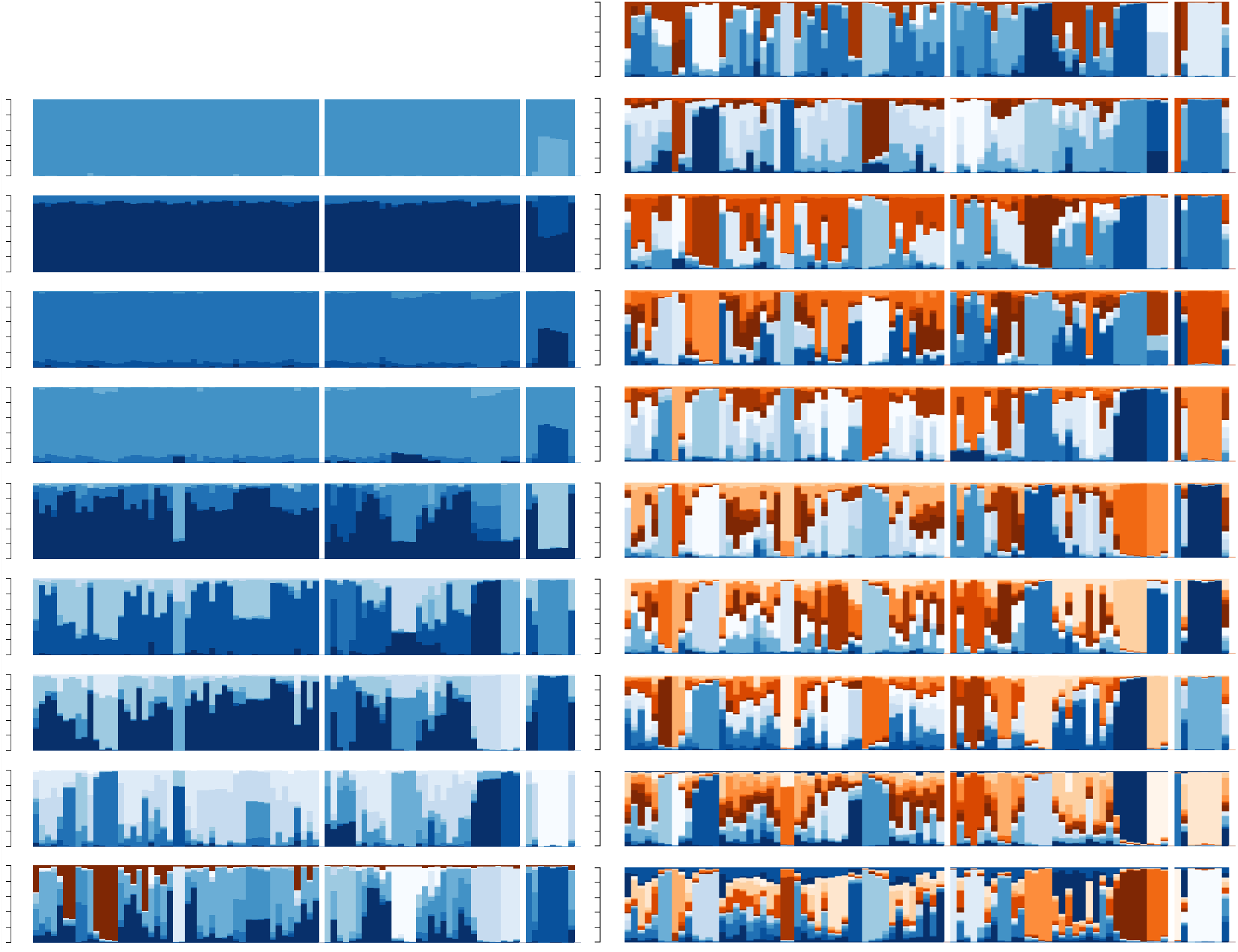
STRUCTURE results for the Core Range GBS data set. On the left are the results for K=2 (top) to K=10 (bottom); on the right are the results for K=11 to K=20. The thin vertical bars represent individual milkweeds, and the three populations (left-to-right northeast, southeast, and European) are separated by thin white bars. Each bar is colored according to the cluster(s) to which it belongs. The optimal model was, K=5, which shows a distinct cluster in Europe, and all US milkweeds largely belonging to the same cluster.

**Figure SI4:**
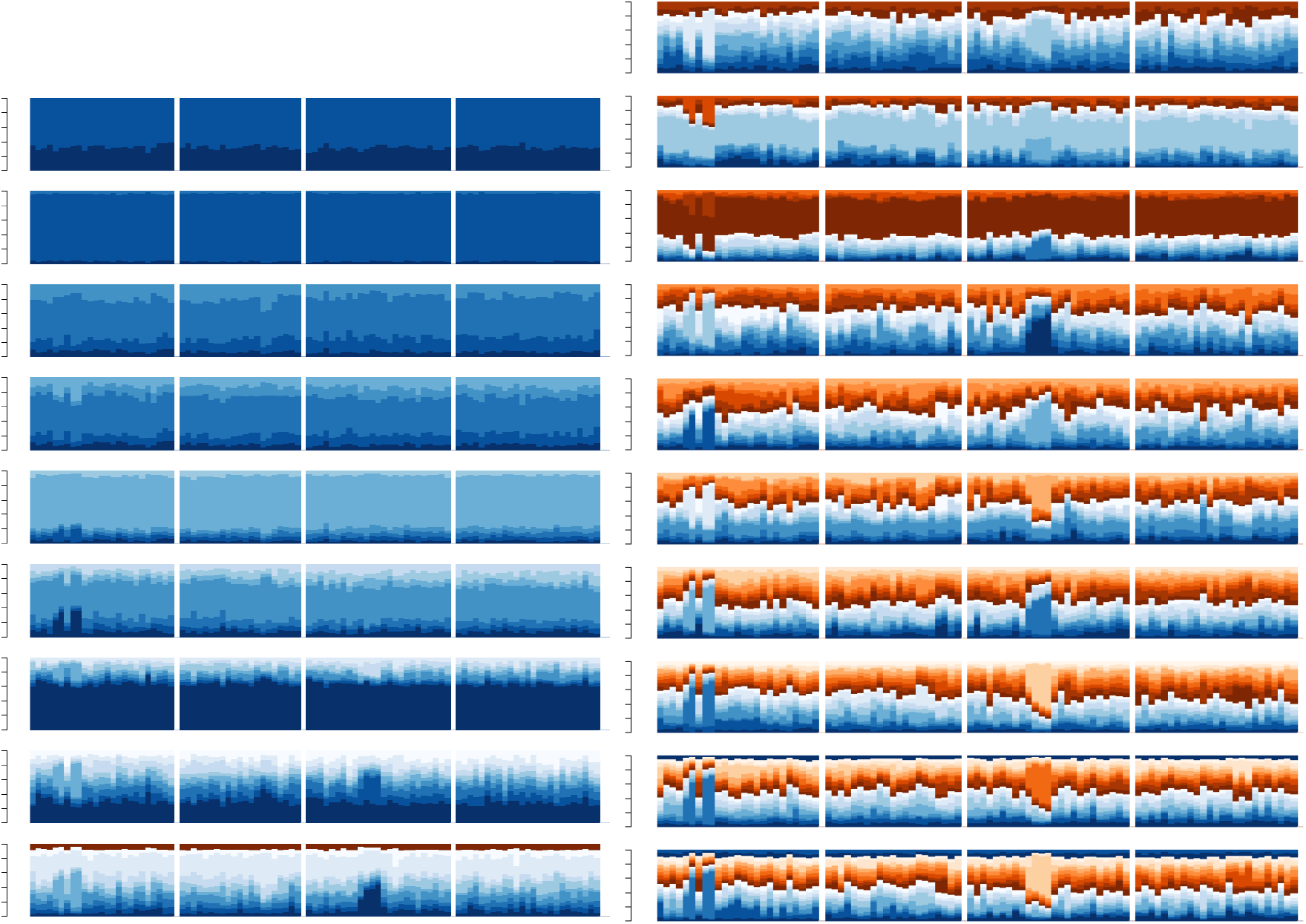
STRUCTURE results for the Broad Range GBS data set. On the left are the results for K=2 (top) to K=10 (bottom); on the right are the results for K=11 to K=20. The thin vertical bars represent individual milkweeds, and the four populations (left-to-right northwest, southwest, northeast, and southeast) are separated by thin white bars. Each bar is colored according to the cluster(s) to which it belongs. Regardless of K-value, the ancestry of milkweeds in this data set is nearly homogenous across the range.

**Figure SI5:**
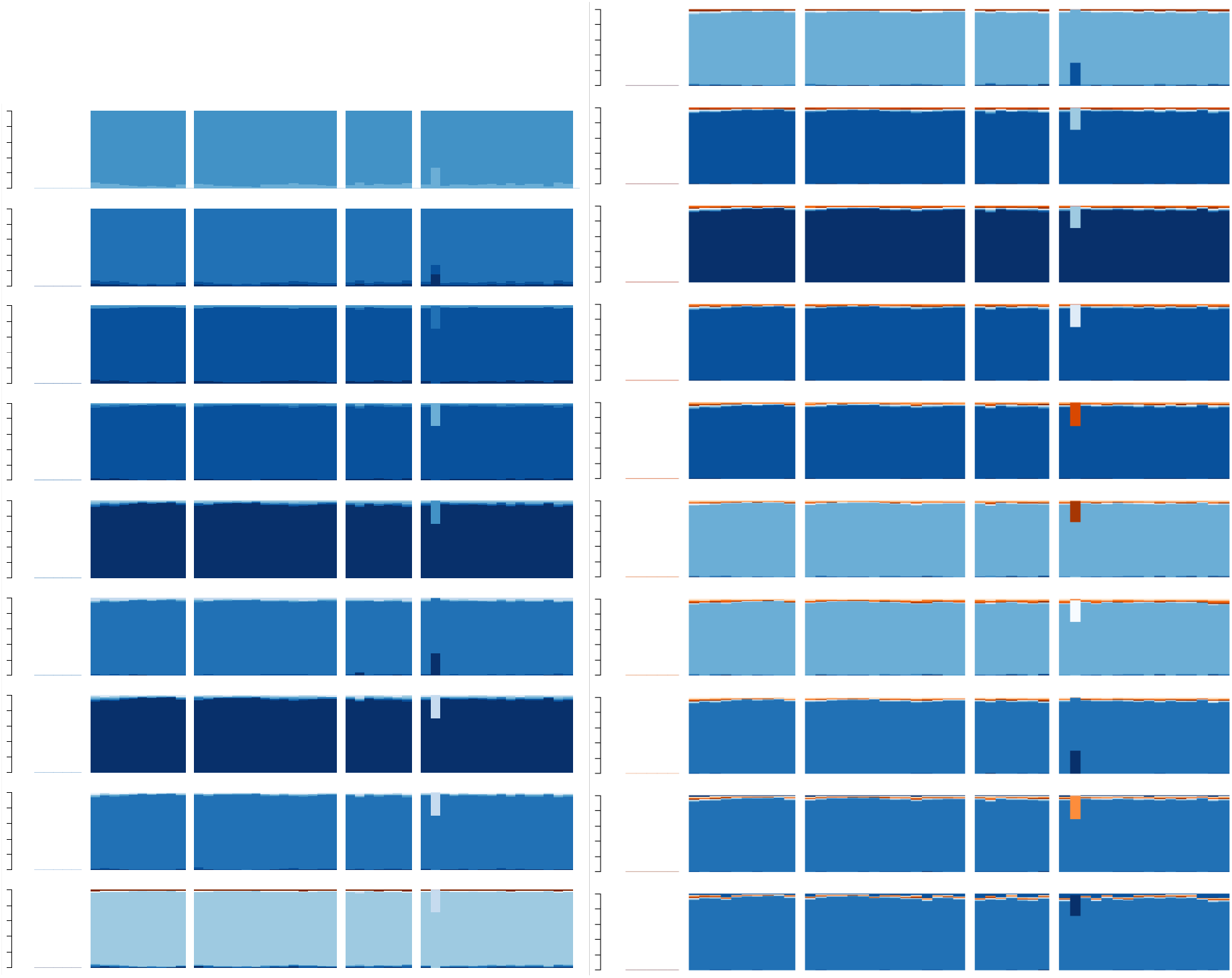
STRUCTURE results for the Broad Range WGR data set. On the left are the results for K=2 (top) to K=10 (bottom); on the right are the results for K=11 to K=20. The thin vertical bars represent individual milkweeds, and the four populations (left-to-right northwest, southwest, northeast, and southeast) are separated by thin white bars. Each bar is colored according to the cluster(s) to which it belongs. Regardless of K-value, the ancestry of milkweeds in this data set is nearly homogenous across the range.

**Figure SI5.5:**
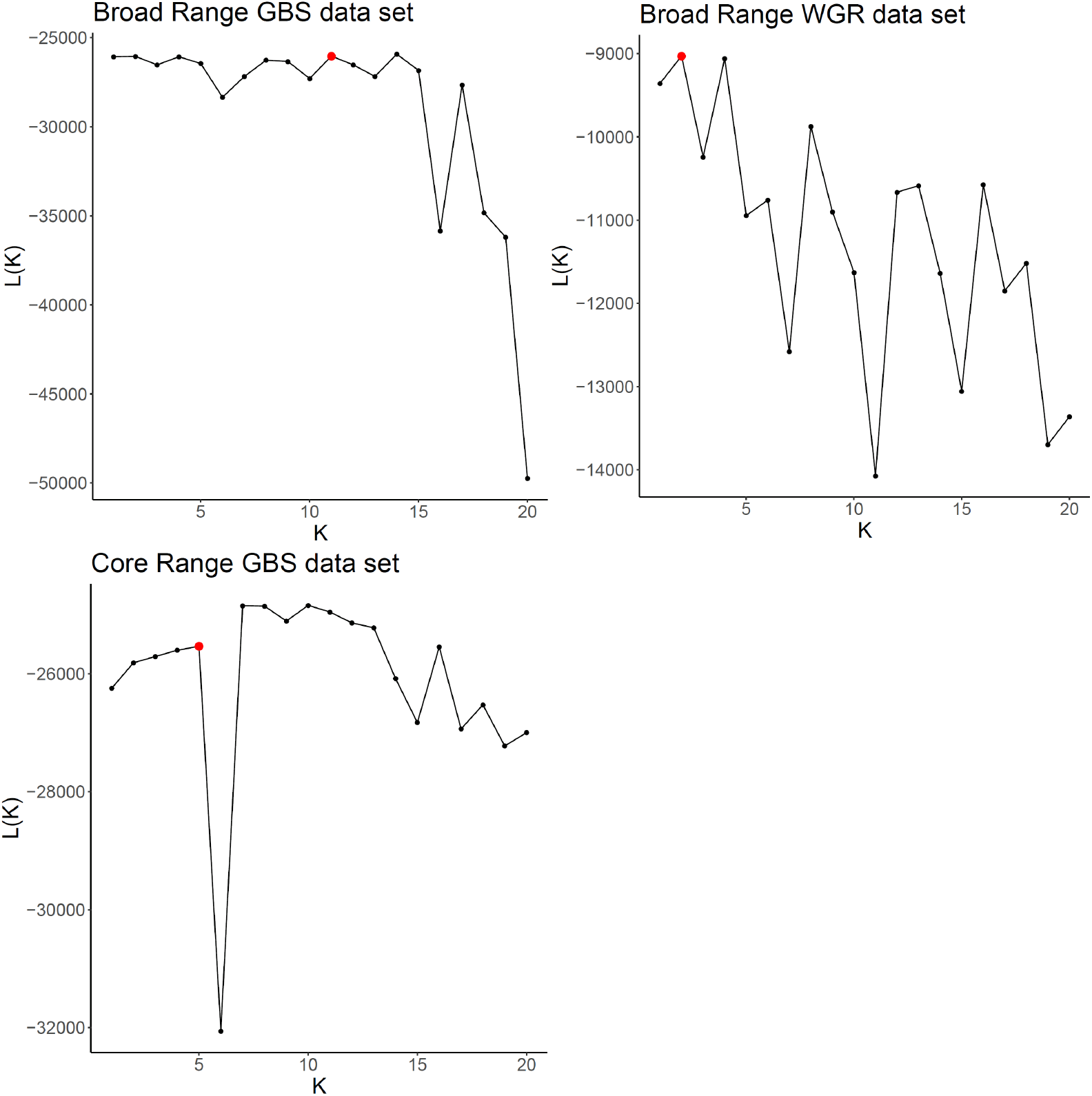
STRUCTURE-produced likelihoods of our data sets under different K-values. Likelihoods of our data (y-axis) under each K-value scenario (x-axis) are shown here. In all three data sets, there is no strong improvement in likelihood from increasing K-values past K=1. The K-value shown in red is the one chosen by the Evanno method; however, as we discuss in the text, the Evanno method is unable to select K=1, which we consider to best capture the data due to other lines of evidence.

**Table 1.1:** Population genetics of A. syriaca and D. plexippus.

| Data Set | SNPs | Missing data | Population | n | H<sub>o</sub> | H<sub>e</sub> | F<sub>IS</sub> | F<sub>ST</sub> |
| --- | --- | --- | --- | --- | --- | --- | --- | --- |
| Broad Range GBS | 891 | 5.2% | Total | 96 | 0.074 | 0.087 | 0.147 | 0.008(1) |
|  |  |  | Northwest | 25 | 0.076 | 0.089 | 0.107 |  |
|  |  |  | Southwest | 21 | 0.077 | 0.087 | 0.085 |  |
|  |  |  | Northeast | 25 | 0.071 | 0.085 | 0.120 |  |
|  |  |  | Southeast | 25 | 0.073 | 0.087 | 0.103 |  |
| Broad Range WGR | 885 | 3.4% | Total | 48 | 0.039 | 0.050 | 0.222 | - 0.002(2) |
|  |  |  | Northwest | 6 | 0.044 | 0.053 | 0.102 |  |
|  |  |  | Southwest | 16 | 0.041 | 0.059 | 0.242 |  |
|  |  |  | Northeast | 10 | 0.035 | 0.045 | 0.144 |  |
|  |  |  | Southeast | 16 | 0.036 | 0.044 | 0.132 |  |
| Core Range GBS | 926 | 4.3% | Total | 87 | 0.076 | 0.088 | 0.134 | 0.039(1) |
|  |  |  | Northeast | 47 | 0.081 | 0.080 | 0.083 |  |
|  |  |  | Southeast | 32 | 0.085 | 0.092 | 0.063 |  |
|  |  |  | Europe | 8 | 0.062 | 0.080 | 0.176 |  |
| Monarch | 11,703 | 3.6% | Total | 28 | 0.109 | 0.124 | 0.125 | n/a |

1: AMOVA, $p < 1 \times 10^{-4}$
2: AMOVA, $p = 0.47$

**Table 1.2:** Population structure of A. syriaca.

| Data Set | Pairwise comparison | Pairwise $F_{ST}$ (GBS) | Pairwise $F_{ST}$ (WGR) |
| --- | --- | --- | --- |
| Broad Range | Northwest vs Southwest | 0.009 | -0.021 |
|  | Northwest vs Northeast | 0.019 | 0.004 |
|  | Northwest vs Southeast | 0.017 | -0.000 |
|  | Southwest vs Southeast | 0.009 | 0.002 |
|  | Southwest vs Northeast | 0.011 | 0.008 |
|  | Northeast vs Southeast | 0.002 | -0.000 |
| Core Range | Northeast vs Southeast | 0.009 |  |
|  | Northeast vs Europe | 0.082 |  |
|  | Southeast vs Europe | 0.081 |  |

GBS: Data from Genotyping By Sequencing approach. WGR: Data from Whole Genome Resequencing approach.

#### Population Genetic Analysis

##### F_ST_ analysis and basic population genetic statistics

Population genetic statistics for each of the populations are shown in Tables 1.1 and 1.2. The genetic differentiation of the subpopulations was low, but statistically significant for the GBS data sets (F_ST_ = 0.008 for Broad Range; 0.052 for Core Range; AMOVA *p* < 1*10^-4^ for both). For the Broad Range WGR data set, genetic differentiation was even lower, and not significant (Fst = -0.002, or effectively zero, AMOVA *p* = 0.47), possibly due to the smaller number of individuals in each population. In the Core Range GBS data set, the greatest pairwise F_ST_ was between the invasive European population and native populations; pairwise F_ST_ was lower between the northeast and southeast populations by a factor of 10. In the Broad Range GBS data set, the greatest pairwise F_ST_ was between the Northwest population and the two eastern populations, although even this was relatively low, at 0.02. Within each dataset, heterozygosity was relatively constant among populations, with the exception that both observed and expected heterozygosity were lower in Europe than in the other populations in the Core Range data set, showing reduced genetic diversity in the invasive range of *A. syriaca.* The *A. syriaca* specimen chosen for genome sequencing was an invasive, European milkweed, on the logic that the invasion process had likely led to more inbreeding than is usual in other *A. syriaca* populations, and the reduced heterozygosity of this population suggests that this was indeed the case. The reduced heterozygosity is beneficial for genome assembly.

##### STRUCTURE analysis

Applying the Evanno method to our STRUCTURE results resulted in an optimal number of *k* = 5 (Figure C) for the Core Range Data Set. Examination of the STRUCTURE results shows a very similar pattern for all values between *k* = 2 and *k* = 5: a single cluster dominates all individuals from North America, and a second cluster is found in a number of invasive *A. syriaca* collected from Europe (Figure SI3). Other clusters, when present, account for very little of the ancestry of any *A. syriaca* specimens. For the Broad Range data sets, the Evanno method selected *k = 11* for the GBS data set and *k* = 2 for the WGR data set. However, the Evanno method is unable to consider *k = 1* as the best cluster, since it uses changes in the likelihood of the data between *k = x* and *k = x-1.* Visualizing the cluster results showed patterns in which each genetic cluster was found in every individual to a similar extent, which suggests that there is minimal geographic structuring within the Broad Range data set.

##### PCA analysis

**Figure SI6:**
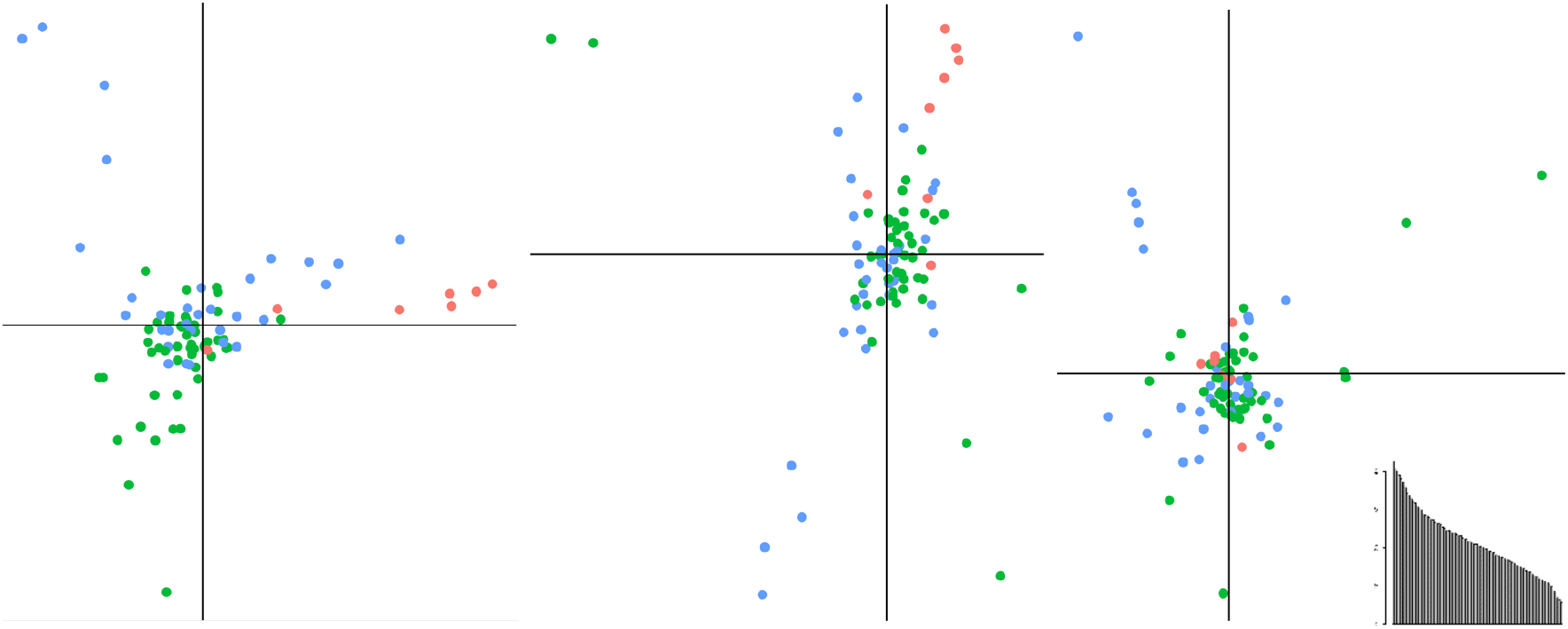
PCA plots for the Core Range GBS data set. PCA plots of the first six PC axes (left: PC1 on the x-axis, PC2 on the y-axis; middle: PC3 and PC4; right: PC5 and PC6). Points are colored according to their population: green is northeast, blue is southeast, red is Europe. Eigenvalues are show in the inset.

**Figure SI7:**
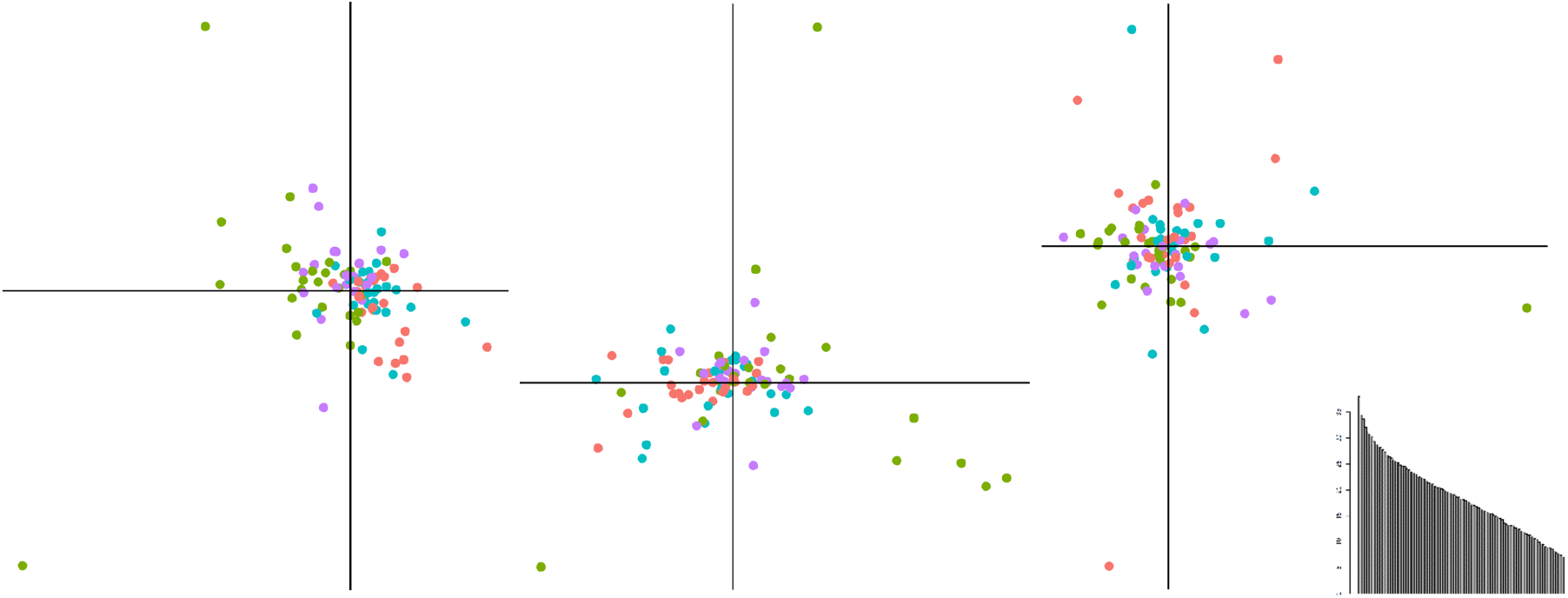
PCA plots for the Broad Range GBS data set. PCA plots of the first six PC axes (left: PC1 on the x-axis, PC2 on the y-axis; middle: PC3 and PC4; right: PC5 and PC6). Points are colored according to their population: green is northwest, purple is southwest, red is northeast, and teal is southeast. Eigenvalues are show in the inset.

**Figure SI8:**
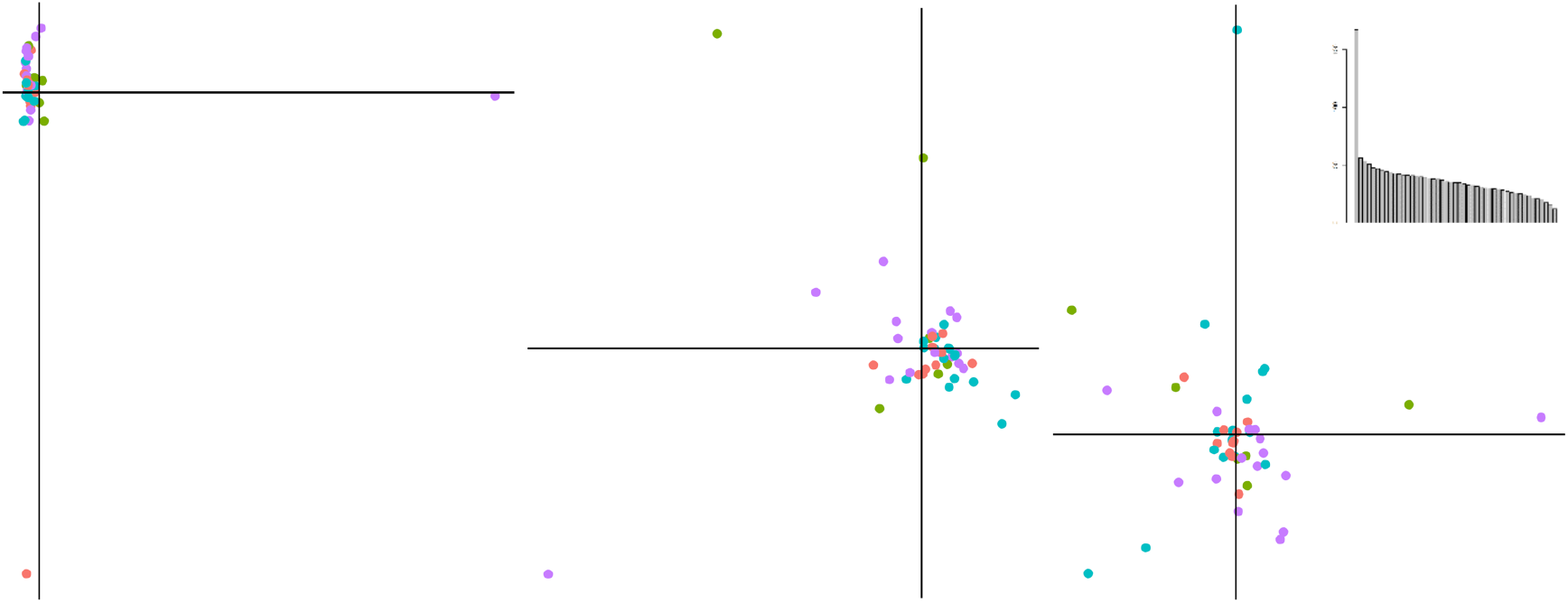
PCA plots for the Broad Range WGR data set. PCA plots of the first six PC axes (left: PC1 on the x-axis, PC2 on the y-axis; middle: PC3 and PC4; right: PC5 and PC6). Points are colored according to their population: green is northwest, purple is southwest, red is northeast, and teal is southeast. Eigenvalues are show in the inset.

For all three data sets, none of the first six PC axes clearly separate any population from any other(s); although some PC axes show some degree of geographic structure, there is always a considerable degree of overlap between the PC values of the various populations.

For the Core Range GBS data set (Figure SI6), PC1 distinguishes several individuals from the European population from the North American population, while PC2, and to a lesser extent PC3, show a limited degree of separation between northern and southern populations.

For the Wide Range GBS data set (Figure SI7), PC1 largely separates several northwestern individuals from the remainder of the data set, possibly indicating introgression from *A. speciosa*, which is known to hybridize with *A. syriaca* in the northwestern part of the *A. syriaca* range, and PC2 and PC3 somewhat distinguish western and eastern populations, but the other PC axes show very little geographic patterning.

For the Wide Range WGR data set (Figure SI8), PC1 and PC2 separate a single individual each from the remainder of the individuals in the data set. Little geographic signal is visible in the remaining PC axes.

The insets show the eigenvalues for each principal component; these decline quite slowly, indicating that each individual PC axis explains relatively little of the variation in genotype. The exception is PC1 of the Broad Range WGR data set, which distinguishes one southwestern individual from the remaining milkweeds, perhaps also representing introgression from another species.

#### Demographic Modelling

Projecting our observed data onto the LDA axes of our simulated data indicated that our set of demographic models were realistic, as the observed data fell within or near the cloud of simulated data points along all LDA axes (Figure SI9).

**Figure SI9:**
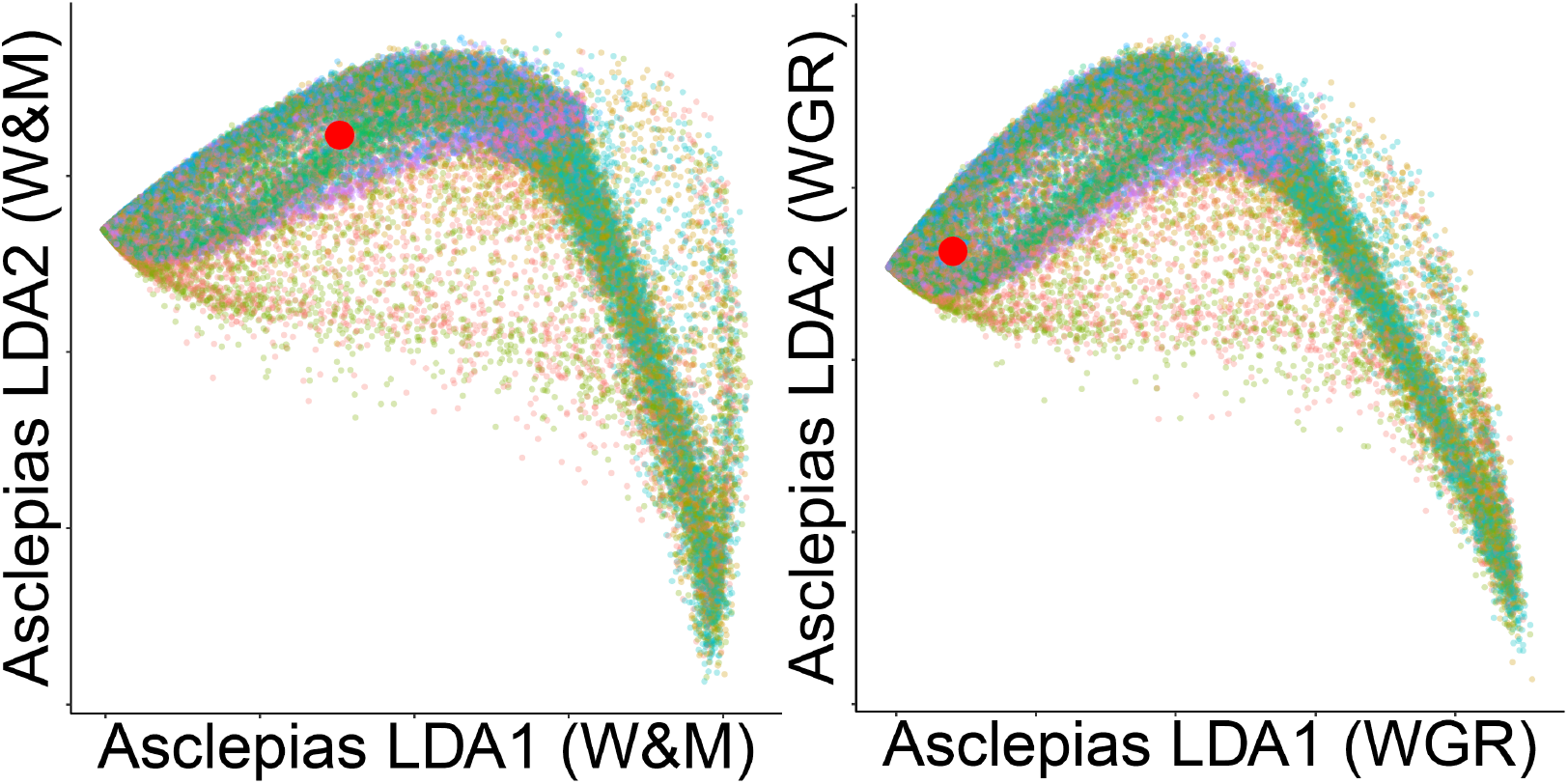
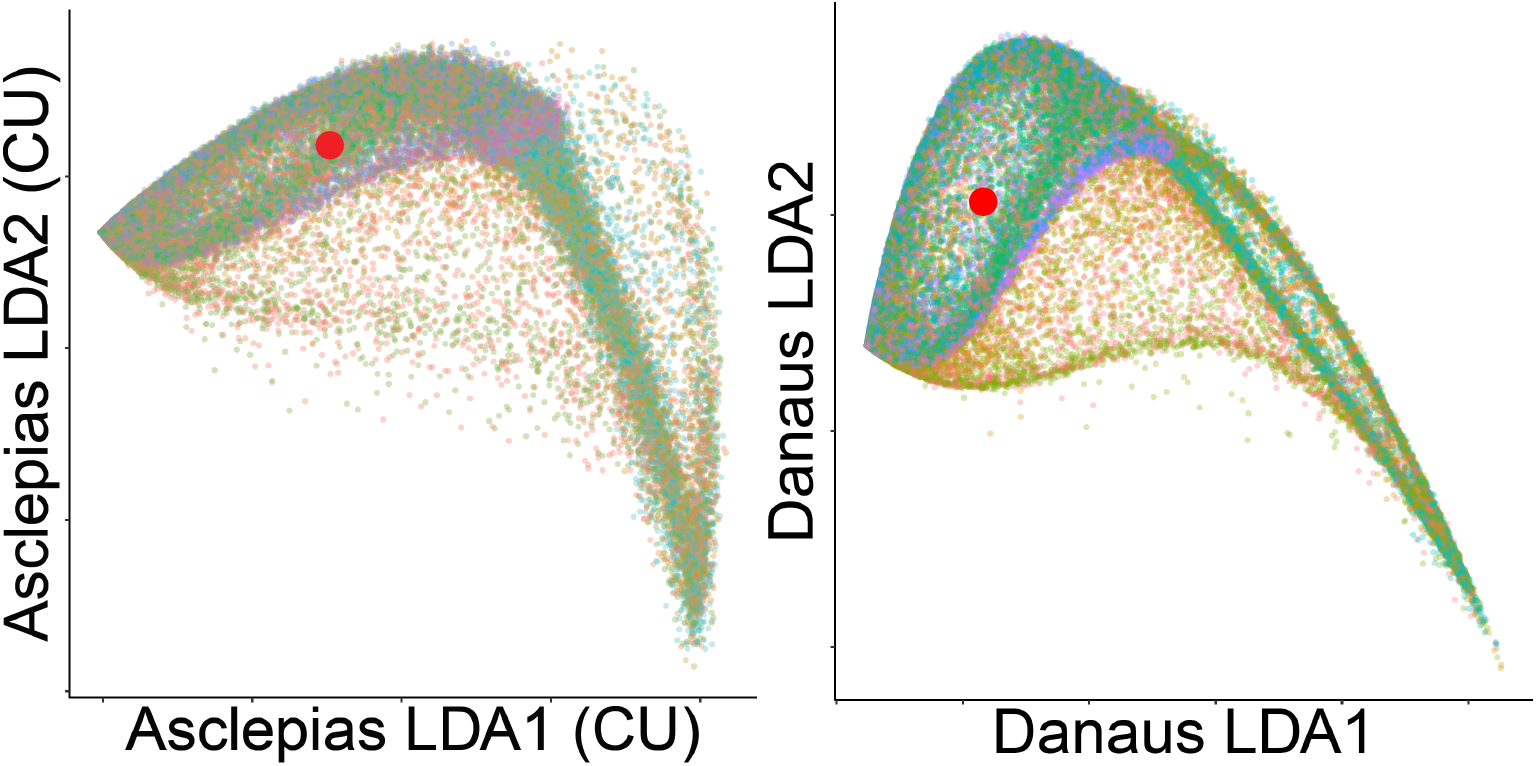
Our simulations captured the characteristics of our observed data set. LDA plots for *Asclepias syriaca* data sets (upper left: Broad Range GBS; upper right: Broad Range WGR; lower left: Core Range WGR), and for *D. plexippus* (lower right). Small points represent simulated data sets, colored according to the demographic model used to simulate them. The large red point represents our observed data set.

Per Pudlo *et al*. (2016), we also confirmed that we produced enough simulations, as the prior error rate decreased only slightly by the addition of the last 20% of simulations (table SI3 below). In fact, we found a few cases in which error rates went up slightly after adding the final 20% of the data (by 0.3% or less), indicating that we are in the regime in which changes in error rate are determined by random fluctuations, and confirming that adding more simulations will not further improve the accuracy of this method.

**Table SI3:** 10,000 simulations per demographic scenario was sufficient.

|  | Prior Error Rates |  |  |  |  |  |  |  |
| --- | --- | --- | --- | --- | --- | --- | --- | --- |
| Data set: | *A. syriaca* (W&M GBS) |  | *A. syriaca* (W&M WGR) |  | *A. syriaca* (Cornell) |  | *D. plexippus* |  |
| Simulations used | 80% | 100% | 80% | 100% | 80% | 100% | 80% | 100% |
| Identifying demographic model | 74.04 % | 74.05 % | 75.18 % | 75.32 % | 74.62 % | 74.59 % | 69.22 % | 69.32 % |
| Identifying presence/absence of recent bottleneck | 38.92 % | 38.84 % | 39.56 % | 39.76 % | 38.72% | 38.75 % | 31.50 % | 31.55 % |
| Identifying presence/absence of recent expansion | 38.05 % | 38.17 % | 38.59 % | 38.33 % | 38.42% | 38.39 % | 34.27 % | 34.36 % |
| Identifying presence/absence of ancient expansion | 28.22 % | 28.08 % | 28.65 % | 28.95 % | 28.59% | 28.41 % | 29.82 % | 29.78 % |

Finally, we followed the recommendation of Pudlo et al. (2016) for determining whether we had used enough decision trees in our Random Forest algorithm. To do this, we repeated the RF algorithm several times using fewer trees, recalculating the prior error rate each time. If the error rate stays nearly flat as we approach the maximum number of trees, this means that we used an appropriate number of trees, which was indeed the case for all three data sets (Figure SI10).

**Figure SI10:**
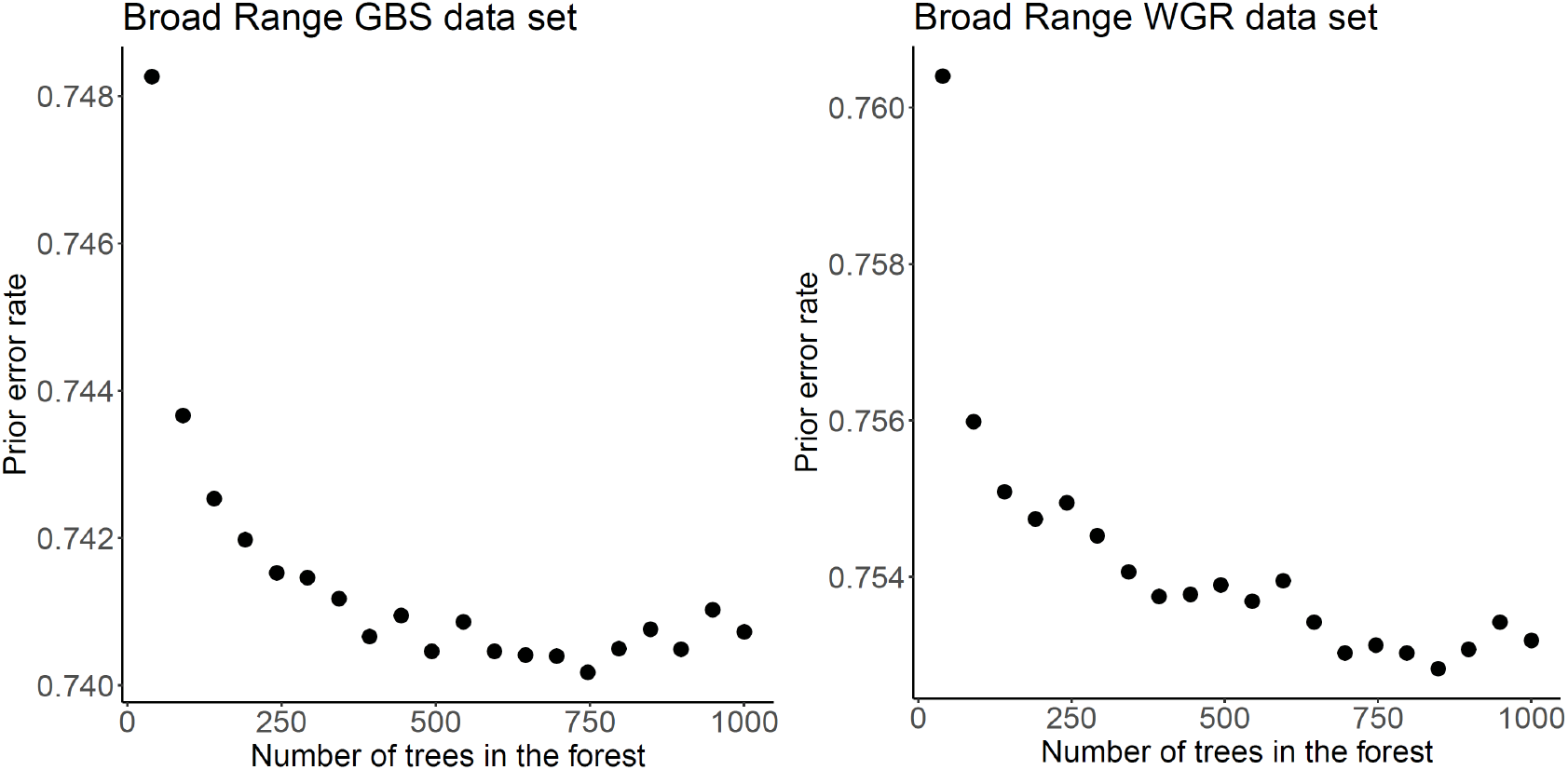
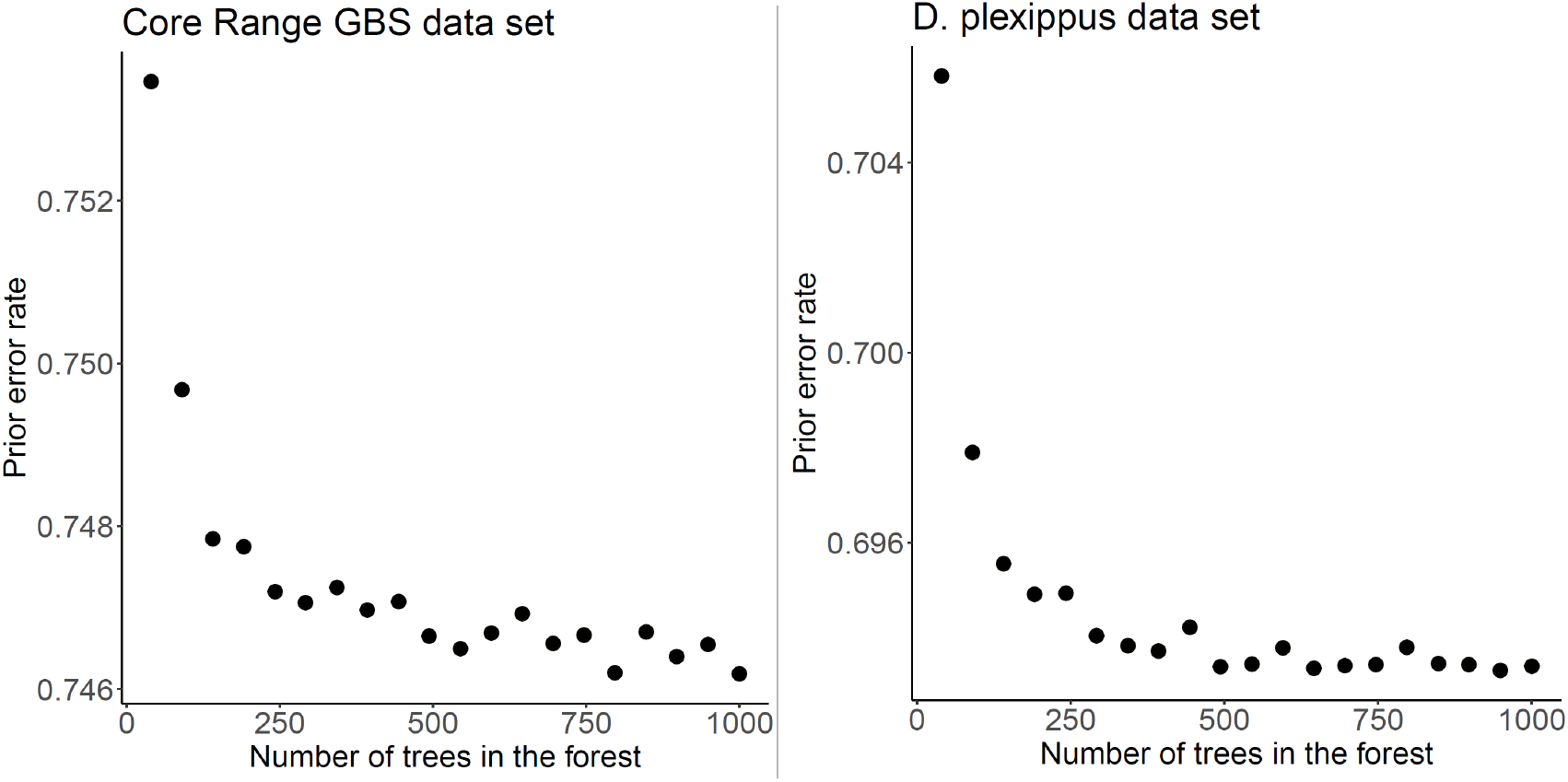
1000 trees in the Random Forest was a sufficient number.

The single best model for milkweeds differed slightly among our three datasets. For the Broad Range GBS data set, the single best model included both ancient and century-scale expansions, but no recent bottleneck; however, this model had a posterior probability of only 0.15. For the Broad Range WGR data set, the single best model was the one that included both the two expansions, but also the bottleneck (0.28). For the Core Range data set, the single best model was the same as for the Broad Range GBS data set, but with higher posterior probability (0.85), including recent expansion and ancient expansion, but no recent bottleneck. For the monarch data set, the single best model includes a recent and ancient expansion, but no recent bottleneck (0.67 posterior probability), the same model chosen in both *A. syriaca* GBS data sets. Due to the fairly high prior error rates when estimating individual models, we focus our attention on estimating the presence or absence of each demographic event separately; these results are described in the main text.

Model parameters estimated with the ABC-RF approach were nearly identical to their prior distributions, suggesting that our dataset does not have sufficient resolution for parameter estimation.

**Table 2:**
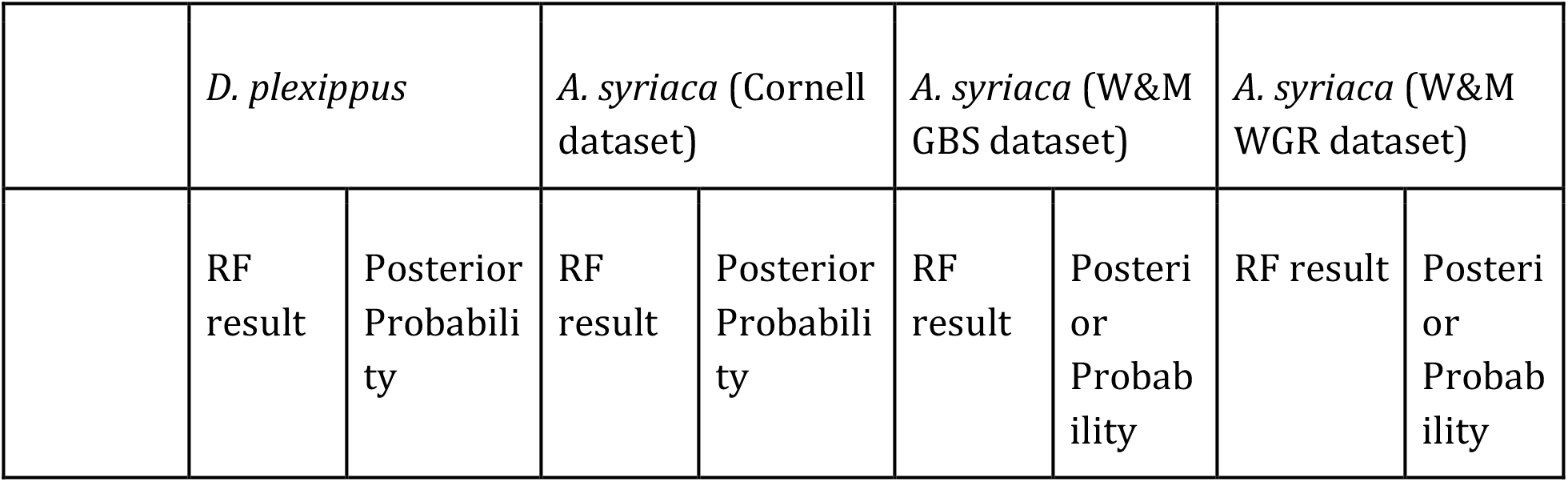
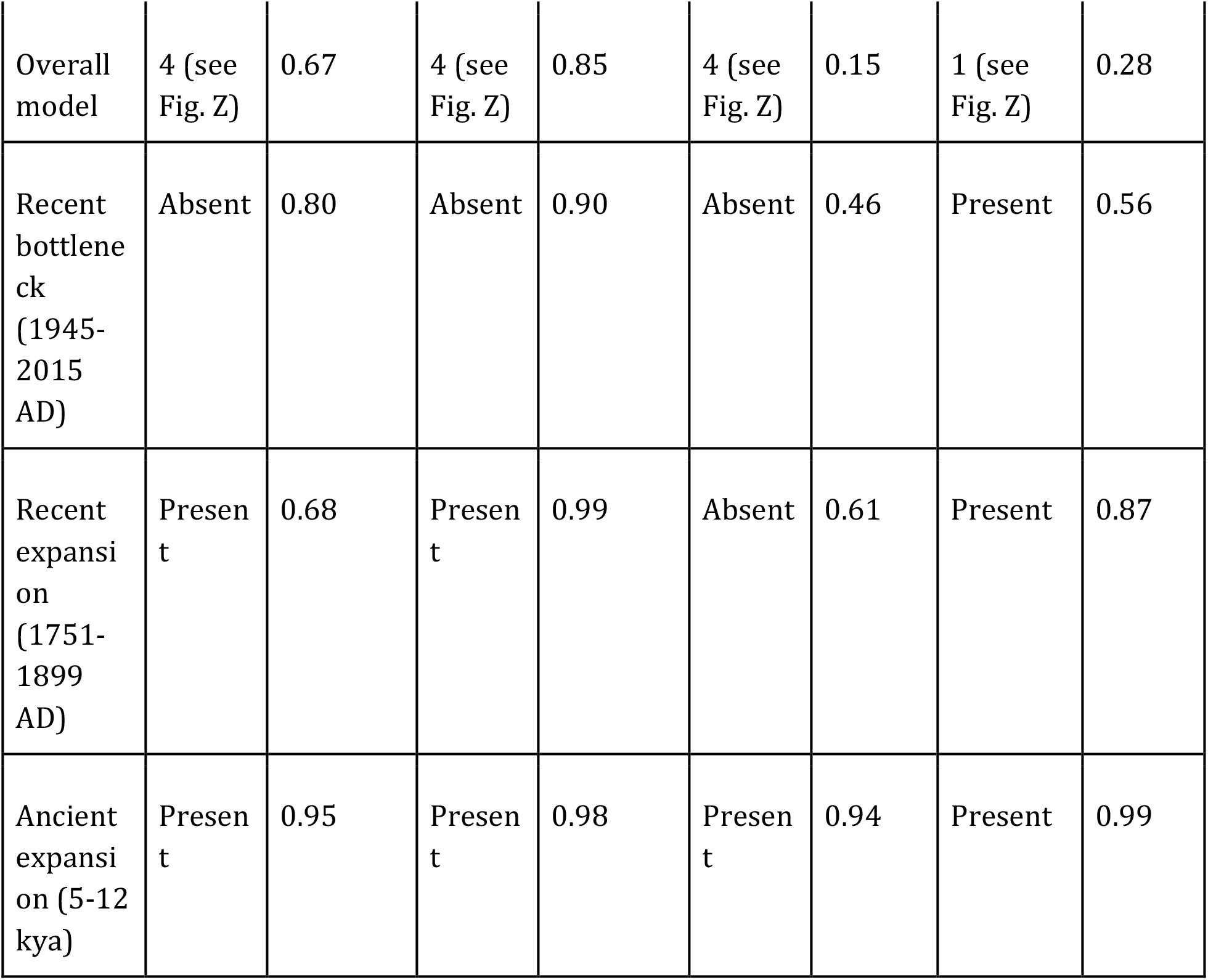
Model selection by ABC-RF.

**Figure 3.**
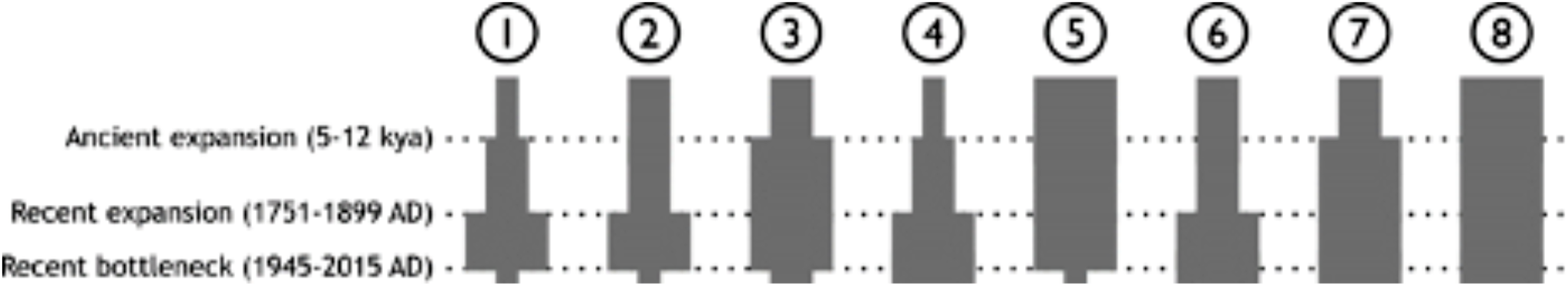
(ABC models. Summary of results)

### Supplementary File SA

We used the following D. plexippus libraries from Zhan et al. (2014):

| NCBI Library | Collection locality |
| --- | --- |
| SRR1548504 | MA, USA |
| SRR1548506 | MA, USA |
| SRR1548571 | MA, USA |
| SRR1548575 | MA, USA |
| SRR1548576 | MA, USA |
| SRR1548577 | MA, USA |
| SRR1548578 | MA, USA |
| SRR1548572 | NJ, USA |
| SRR1548573 | NJ, USA |
| SRR1548574 | NJ, USA |
| SRR1551995 | CA, USA |
| SRR1552224 | CA, USA |
| SRR1552225 | CA, USA |
| SRR1552204 | Mexico |
| SRR1552205 | Mexico |
| SRR1552206 | Mexico |
| SRR1552207 | Mexico |
| SRR1552208 | Mexico |
| SRR1552209 | Mexico |
| SRR1549524 | North FL, USA |
| SRR1549525 | North FL, USA |
| SRR1549526 | North FL, USA |
| SRR1552001 | North FL, USA |
| SRR1552222 | North FL, USA |
| SRR1552223 | North FL, USA |
| SRR1549527 | TX, USA |
| SRR1549528 | TX, USA |
| SRR1549529 | TX, USA |
| SRR1980588 | TX, USA |

### Supplementary File 2: Identifying Batch Effects

We identified SNPs from the combined Cornell and W&M datasets using the same stacks pipeline described in the main text. This resulted in 872 SNP markers from 181 *A. syriaca* individuals. These markers were then used in a STRUCTURE analysis identical to that described in the main text, with the exception that we only analyzed possible numbers of clusters between *K* = 2 and *K* = 10.

STRUCTURE results were processed and visualized using the same pipeline described in the main text. The results are shown below in Figure S2.1.

For many values of *K*, the differences between the STRUCTURE results for the Cornell data set and the W&M data set were subtle: for instance, for *K* = 2, Cornell individuals had approximately 25-35% ancestry from Cluster 1, while W&M individuals had around 35-35% ancestry the same cluster (Figure S2.1). We therefore also used a second clustering method implemented in the adegenet 2.1.2 package (Jombart 2008, Jombart and Ahmed 2011) in R, which uses a *K*-means approach to assign individuals to one of *K* clusters, with the appropriate *K* chosen based on the Bayesian Information Criterion.

Runs with *K* = 2 and *K* = 3 produced the two lowest BICs, which were nearly equal. Both runs produced similar results, with the cluster assignments almost exactly mirroring membership in the Cornell or W&M datasets (Table S2.1). The difference between the two is that at *K* = 3, some European individuals from the Cornell data set were split off from the remainder of the Cornell individuals (results which were also seen in the STRUCTURE results (Figure S2.1).

**Table S2.1:** Adegenet assigns individuals from different data sets to different clusters.

| Cluster Assignment | W&M individuals | Cornell individuals |
| --- | --- | --- |
| $K = 2$, cluster 1 | 90 | 1 |
| $K = 2$, cluster 2 | 4 | 86 |
| $K = 3$, cluster 1 | 5 | 82 |
| $K = 3$, cluster 2 | 0 | 5 |
| $K = 3$, cluster 3 | 89 | 0 |

**Figure 2.1:**
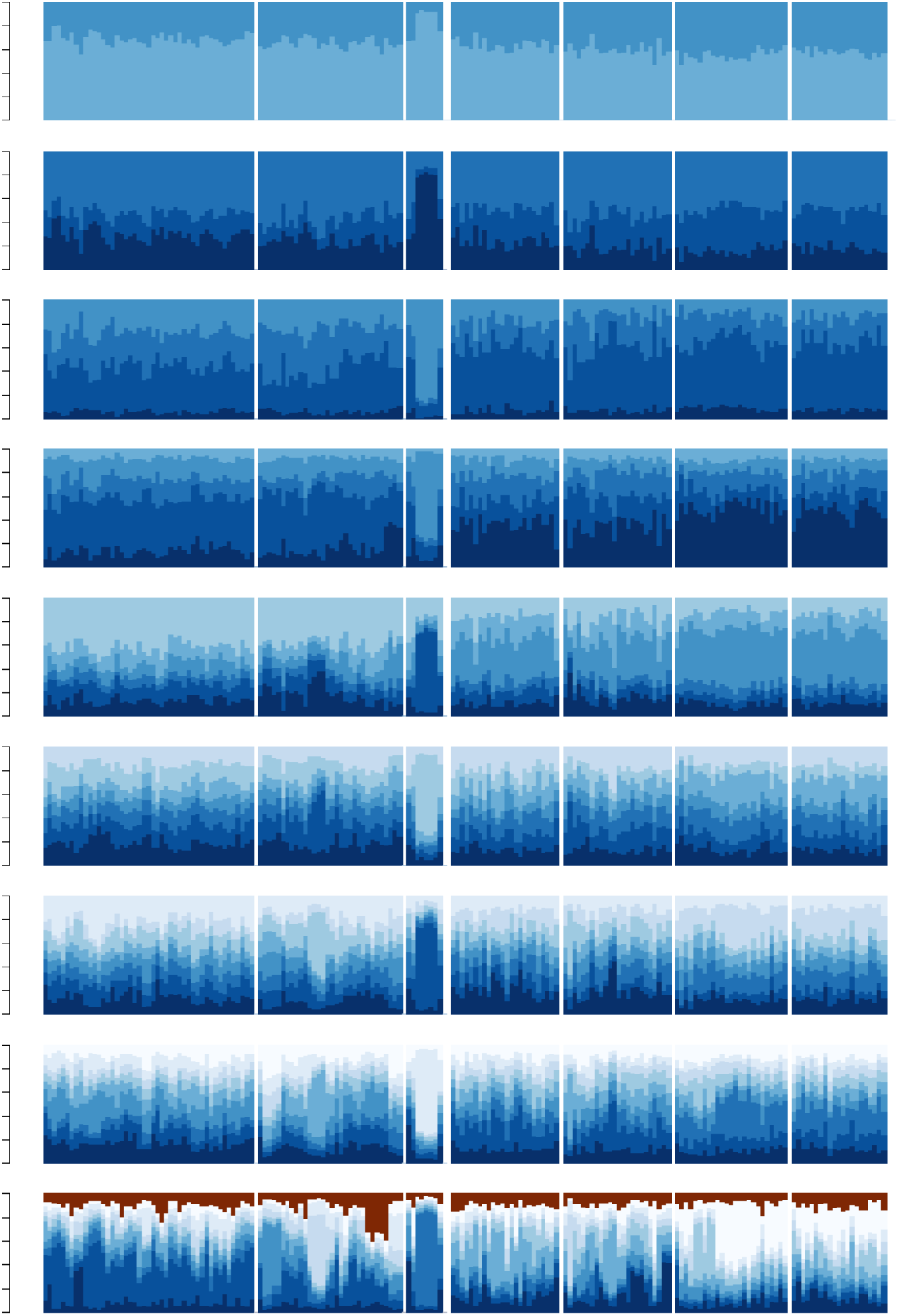
Batch effects appear when attempting to combine Cornell and W&M data sets. The thin vertical bars represent individual milkweeds, divided into seven blocks, from left to right: Cornell Northeast, Cornell Southeast, Cornell Europe, W&M Northeast, W&M Southeast, W&M Northwest, W&M Southwest. Each bar is colored according to the cluster(s) to which it belongs. The top graph shows the results when Structure assumes 2 clusters, then 3 clusters, etc, with the bottommost graph showing *k*=10 clusters. Several of the European individuals form a distinct cluster for all values of *k.* Note the differences in which clusters are common in North American milkweeds in the Cornell data set vs the W&M data set.

